# Glutamatergic neuronal activity regulates angiogenesis and blood-retinal barrier maturation via Norrin/β-catenin signaling

**DOI:** 10.1101/2023.07.10.548410

**Authors:** Saptarshi Biswas, Sanjid Shahriar, Galina Bachay, Panos Arvanitis, Danny Jamoul, William J. Brunken, Dritan Agalliu

## Abstract

Interactions among neuronal, glial and vascular components are crucial for retinal angiogenesis and blood-retinal barrier (BRB) maturation. Although synaptic dysfunction precedes vascular abnormalities in many retinal pathologies, how neuronal activity, specifically glutamatergic activity, regulates retinal angiogenesis and BRB maturation remains unclear. Using *in vivo* genetic studies in mice, single-cell RNA-sequencing and functional validation, we show that deep plexus angiogenesis and paracellular BRB maturation are delayed in *Vglut1*^-/-^ retinas where neurons fail to release glutamate. In contrast, deep plexus angiogenesis and paracellular BRB maturation are accelerated in *Gnat1*^-/-^ retinas where constitutively depolarized rods release excessive glutamate. Norrin expression and endothelial Norrin/β-catenin signaling are downregulated in *Vglut1*^-/-^ retinas, and upregulated in *Gnat1*^-/-^ retinas. Pharmacological activation of endothelial Norrin/β-catenin signaling in *Vglut1*^-/-^ retinas rescued defects in deep plexus angiogenesis and paracellular BRB maturation. Our findings demonstrate that glutamatergic neuronal activity regulates retinal angiogenesis and BRB maturation by modulating endothelial Norrin/β-catenin signaling.

## INTRODUCTION

The crosstalk among neuronal, glial, and vascular components is essential, not only for proper angiogenesis, but also for the establishment of the inner blood-retinal barrier (BRB) in the developing retina ^1^. Retinal angiogenesis is tightly coordinated with the maturation of the BRB properties by endothelial cells (ECs), through the presence of tight-junctions (TJ) that limit paracellular permeability between ECs, specialized transporters to facilitate bidirectional transport of nutrients, and suppression of transcytosis that reduces transcellular EC permeability ^2, 3^. Although multiple signaling pathways operating in ECs, pericytes (PCs), neuronal or glial cells have emerged as critical regulators of retinal angiogenesis and BRB maturation [reviewed in ^4^], it is unclear how distinct types of neuronal synaptic activities influence these processes, and whether neurotransmitters act directly on ECs to mediate their effects. This is highly significant since early loss of neurovascular coupling, impaired response of photoreceptors to light, gradual neurodegeneration, gliosis and neuroinflammation are found prior to any obvious vascular pathologies in diabetic retinopathy and other retinal disease ^5–7^. Moreover, it has been postulated that diabetes-induced changes in neurons and glial cells lead to vascular defects ^8^, yet the underlying mechanisms remain elusive.

Distinct types of neuronal activities are present in the postnatal mouse retina during angiogenesis and BRB maturation **(Figure 1A)**. From birth to postnatal day (P) 10, starburst amacrine cells propagate spontaneous cholinergic waves ^9–11^. These cholinergic waves gradually subside and spontaneous glutamatergic waves emerge from P10 to P14 ^11–14^. The majority of neuronal activities in the mouse retina from P14 onwards are mediated by light-dependent glutamatergic synaptic activity of photoreceptors, and photoactivation of intrinsically-photosensitive retinal ganglion cells (ipRGCs) [reviewed in ^4^]. However, light activation of photoreceptors has been observed as early as P8 ^13, 15^. The superficial vascular plexus in the retina develops between P0 - P8 ^16^ over the ganglion cell layer (GCL) synchronous with neuronal activities from the cholinergic waves and photoactivation of ipRGCs. In contrast, the deep plexus angiogenesis (P8 - P12) in the outer plexiform layer (OPL) ^16^ and BRB maturation (P6 - P18) span both cholinergic and glutamatergic spontaneous waves, glutamatergic synaptic activity of photoreceptors as well as photoactivation of ipRGCs [reviewed in ^4^]. Few studies have addressed how neuronal activity regulates retinal angiogenesis and BRB maturation in the developing retina. Photoactivation of both Opn4^+^ ^17^ and Opn5^+^ ^18^ RGCs regulates angiogenesis in the retina. Pharmacological blockade of cholinergic waves before P8 delays deep plexus angiogenesis and BRB maturation, suggesting an important role for cholinergic activity in these processes ^14^. However, the role of glutamatergic neuronal activities in retinal angiogenesis and BRB maturation remains unclear, although it is the predominant type of neuronal activity when most of the vascular development and BRB maturation occurs in the retina.

**Figure 1.**
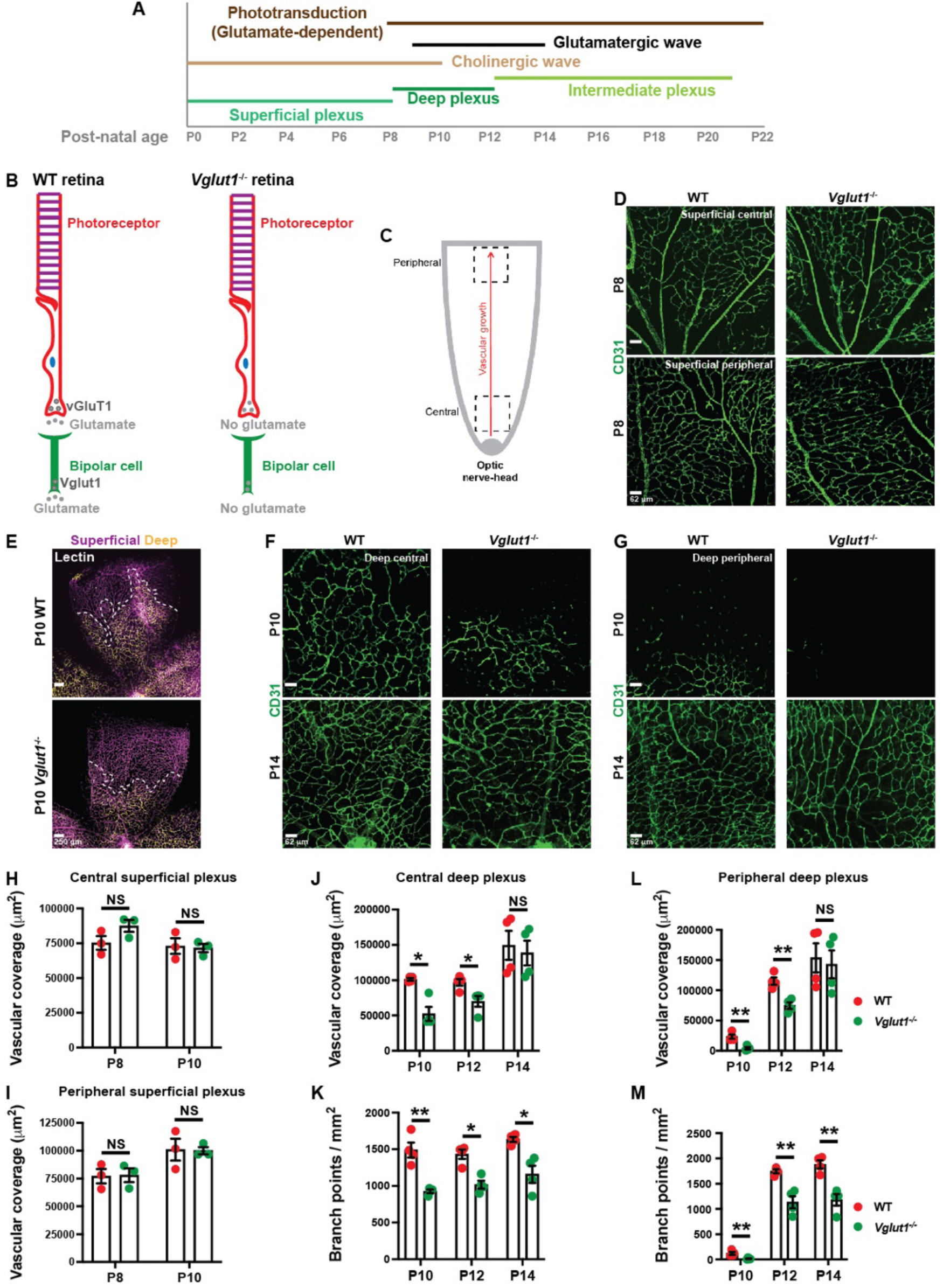
Deep plexus angiogenesis is delayed in the *Vglut1*^-/-^ retina. **A)** Timeline of the neuronal activity [cholinergic and glutamatergic waves and glutamatergic synaptic activity of photoreceptors (phototransduction)] and angiogenesis (superficial, deep and intermediate plexuses) in the postnatal wild-type (WT) retina (P1-P22). **B)** Schematic diagram of glutamate release from bipolar cells and photoreceptors in WT and *Vglut1*^-/-^ retinas. *Vglut1*^-/-^ photoreceptors and bipolar cells fail to release glutamate into the synaptic cleft. **C)** Schematic diagram of distinct developmental regions in the retina. Central retina is developmentally more mature than peripheral retina. **D)** Representative images of the superficial plexus from P8 WT and *Vglut1*^-/-^ retinal flat mounts stained for CD31 (green; EC marker). **E)** Representative images of the retinas from P10 WT and *Vglut1*^-/-^ retinal flat mounts stained for CD31. Superficial plexus is color-coded in purple and deep plexus is color-coded in yellow. Dashed white line shows the extent of deep plexus vascularization. **F, G)** Representative images of the deep plexus from P10 and P14 WT and *Vglut1*^-/-^ retinal flat mounts visualized with CD31 staining (green). **H, I)** Dotted bar graphs of the vascular coverage of the superficial plexus in P8 and P10 WT and *Vglut1*^-/-^ central (**H**) and peripheral (**I**) retina (n = 3 mice/genotype). **J-M)** Dotted bar graphs of the vascular coverage (**J, L**) and branch point density (**K, M**) of the deep plexus in P10, P12 and P14 WT and *Vglut1*^-/-^ central (**J, K**) and peripheral (**L, M**) retina (n = 4 mice/genotype). Scale bars: E = 250 μm; D, F, G = 62 μm. Statistical analyses were done with Students t-test: *p<0.05, **p<0.02, NS = not significant. Error bars: Mean ± S.E.M.

The Norrin/β-catenin signaling is critical for retinal angiogenesis and BRB maturation by inducing stabilization and nuclear translocation of β-catenin. Genetic elimination of Norrin (ligand), Frizzled-4, Lrp5/6, Tspan12 (receptors) and β-catenin in the mouse retina delays superficial vascular plexus angiogenesis, induces loss of the deep plexus ^19–23^ and BRB integrity ^24, 25^. How Norrin expression and endothelial Norrin/β-catenin signaling are related to the glutamatergic activities in the developing retina is also unknown.

Using *in vivo* anatomical and functional characterizations of the vasculature, as well as single-cell transcriptomic analyses and validation studies, we show that deep plexus angiogenesis and paracellular BRB maturation are delayed in *Vglut1*^-/-^ retinas that have reduced synaptic release of glutamate from photoreceptors and bipolar cells ^12^. In contrast, deep plexus angiogenesis is accelerated and paracellular BRB matures precociously in *Gnat1*^-/-^ retinas, where rod photoreceptors remain constitutively depolarized due to the absence of α-Transducin, a Gα protein critical for phototransduction ^26, 27^. These phenotypes are not due to abnormalities in neurovascular unit (NVU) assembly, aberrant specification of neuronal or glial subtypes, or changes in the neuronal metabolic demand. Importantly, *Norrin* mRNA expression in the inner nuclear layer (INL), and as a consequence endothelial Norrin/β-catenin signaling, are downregulated in *Vglut1*^-/-^ retinas, and upregulated in *Gnat1*^-/-^ retinas. Pharmacological activation of endothelial Norrin/β- catenin signaling in *Vglut1*^-/-^ retinas rescues defects in both deep plexus angiogenesis and paracellular BRB integrity. Our study elucidates a critical mechanism by which glutamatergic neuronal activities regulate retinal angiogenesis and BRB maturation by modulating endothelial Norrin/β-catenin signaling in the developing retina. These findings have important implications for development of potential interventions in various neurovascular pathologies.

## RESULTS

### Deep plexus angiogenesis is delayed in the *Vglut1*^-/-^ retina

Photoreceptors release glutamate in the OPL in the dark and suppress glutamate release upon photon capture ^28^. The synaptic release of glutamate is dependent on vGluT1, a vesicular glutamate transporter ^29^. Consequently, *Vglut1*^-/-^ mice have reduced synaptic release of glutamate from photoreceptors and bipolar cells **(Figure 1B)**^12^. To understand if extracellular glutamate levels affect retinal angiogenesis, we compared the vascular growth in the superficial and deep plexuses between wild-type (WT) and *Vglut1*^-/-^ retinas. Since the central retina is developmentally more mature than the peripheral retina, we analyzed these two regions separately **(Figure 1C)**. Although the vascular coverage of the retina in the superficial plexus was similar in both genotypes at P8 and P10 **(Figure 1D, E, H, I)**, the vascular coverage of the retina in the deep plexus and its branch-point density were reduced significantly in the P10 and P12 *Vglut1*^-/-^ retinas **(Figure 1E-G, J-M)**. By P14, however, the vascular coverage of the retina in the deep plexus was similar between the two genotypes, although the branch-point density of the *Vglut1*^-/-^ deep plexus was still significantly lower than the WT **(Figure 1F, G, J-M)**. Importantly, these vascular abnormalities were consistent in both the central and peripheral *Vglut1*^-/-^ retinas (**Figure 1J-M**).

To understand the molecular mechanisms underlying these vascular phenotypes, we performed single-cell RNA-sequencing analyses (scRNA-seq) of ECs that were dissociated from P10 WT and *Vglut1*^-/-^ retinas and sorted for CD31 using fluorescence-activated cell sorting (FACS; **Figure S1A, B)**. The purity of ECs was confirmed by uniform expression of EC markers [*Pecam1* (CD31) and *Cdh5* (VE-Cadherin)] in the entire population, and absence of mural cell [*Pdgfrb* (PDGFRβ) and *Des* (Desmin)] and astrocyte [*Gfap* and *Pdgfra*] markers **(Figure S1C)**. The majority of ECs isolated from either genotype were capillary ECs (Mfsd2a+), as opposed to arterial or venous ECs **(Figure S1D)**, consistent with the predominant distribution of this EC subtype in both the superficial and deep plexuses of the retina (the deep plexus has only capillaries). ECs isolated from P10 WT and *Vglut1*^-/-^ retinas clustered separately in the Uniform Manifold Approximation and Projection (UMAP) plot based on differences in their transcriptomic signatures **(Figure 2A; Tables S1, S2)**. Gene ontology (GO) analyses of the differentially expressed genes (DEGs) between the two genotypes revealed that pathways related to “angiogenesis”, “blood vessel morphogenesis” and “cell proliferation” were downregulated in P10 *Vglut1*^-/-^ compared to WT retinal ECs **(Figure 2B; Tables S3, S5)**, consistent with the phenotypic analyses (**Figure 1**). In contrast, pathways related to “cellular and protein metabolic processes” were upregulated in P10 *Vglut1*^-/-^ compared to WT retinal ECs **(Figure 2C; Tables S4)**. Expression levels of several pro-angiogenic genes [e.g. *Ptprb*, *Aplnr, Unc5b* etc.], including two effectors of Norrin/β-catenin signaling [*Ctnnb1* (β-catenin) and *Lef1*], were downregulated significantly in P10 *Vglut1*^-/-^ compared to WT retinal ECs **(Figure 2D, F; Table S5)**. In contrast, the majority of genes enriched in the deep plexus tip cells (D-tip cells) compared to the superficial plexus tip cells (S-tip cells) [e.g. *Bsg*, *Rgcc*, *Plxnd1*, *Cdkn1a*, *Pde4b*, *Prcp*], as well as those enriched in D-tip cells compared to all ECs [e.g. *Lamb1*, *Pcdh17*, *Adm*] ^30^ were upregulated in P10 *Vglut1*^-/-^ compared to WT retinal ECs **(Figure 2E, G; Table S6)**, indicative of delayed deep plexus angiogenesis at the transcriptome level. To validate these molecular findings, we labeled proliferating cells with Ki67 in both WT and *Vglut1*^-/-^ retinas and assessed whether endothelial stalk cell proliferation was reduced in *Vglut1*^-/-^ retinal ECs (**Figure 2H**) ^31^. Consistent with the scRNA-seq data, there was a significant reduction in the number of proliferating Ki67^+^ ECs in P12 *Vglut1*^-/-^ retinal deep plexus compared to the WT **(Figure 2I, J)**, while the number of apoptotic ECs (cleaved Caspase-3^+^) was not significantly different **(Figure 2J)**. Therefore, the delayed angiogenesis in *Vglut1*^-/-^ retinal deep plexus is due, in part, to reduced EC proliferation. Consistent with the scRNA-seq data, we also observed numerous Dll4^+^ tip cells ^32, 33^ in the leading edge of the *Vglut1*^-/-^ deep plexus at P12, while the WT deep plexus was completely vascularized at this stage **(Figure 2K, L)**. Thus, angiogenesis (EC proliferation) is delayed and the number of endothelial tip cells is increased in the deep, but not superficial, plexus of *Vglut1*^-/-^ retinas lacking extracellular glutamate release by neurons.

**Figure 2.**
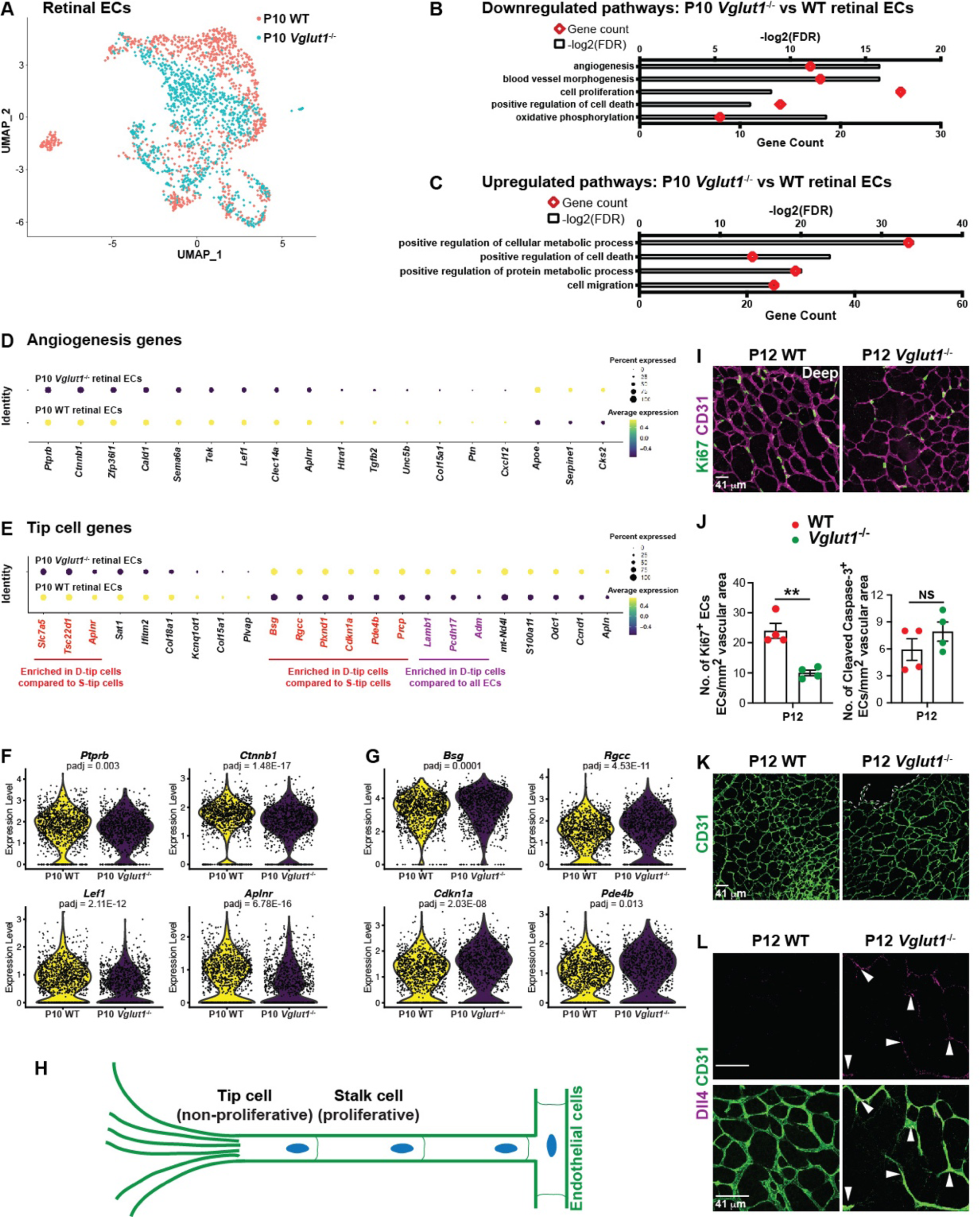
Aberrant angiogenic transcriptomic signatures are present in *Vglut1*^-/-^ ECs from single cell RNA-sequencing (scRNA-seq). **A)** A Uniform Manifold Approximation and Projection (UMAP) for dimension reduction projection shows separation of P10 WT and *Vglut1*^-/-^ retinal ECs based on their transcriptomic signatures. **B, C)** Gene ontology (GO) analyses of P10 WT and *Vglut1*^-/-^ retinal ECs show top downregulated (**B**) and upregulated (**C**) pathways. **D, E)** Dot plot of expression levels for angiogenesis (**D**) and tip cell (**E**) genes in P10 WT and *Vglut1*^-/-^ retinal ECs. The dot size indicates the percent of the population expressing each marker, the color scale indicates the average level of gene expression. Genes enriched in deep plexus tip cells (D-tip cells) compared to superficial plexus tip cells (S-tip cells) are shown in red. Genes enriched in deep plexus tip cells (D-tip cells) compared to all ECs are shown in purple. **F, G)** Violin plots of representative angiogenesis (**F**) and tip cells (**G**) genes in P10 WT and *Vglut1*^-/-^ retinal ECs. **H)** Schematic of a vascular sprout consisting of leading tip cells and proliferative stalk cells. **I)** Representative images of the deep plexus from P12 WT and *Vglut1*^-/-^ retinal flat-mounts stained for Ki67 (green) and CD31 (purple). **J)** Measurements of Ki67^+^ (proliferative), or cleaved Caspase3^+^ (apoptotic) ECs in P12 WT and *Vglut1*^-/-^ retinal deep plexus (n = 4). **K)** Representative images of the deep plexus from P12 WT and *Vglut1*^-/-^ retinal flat-mounts stained for CD31 (green). The dotted line shows the border of the vascular front in the *Vglut1*^-/-^ retina. **L)** Representative images of the deep plexus from P12 WT and *Vglut1*^-/-^ retinal flat-mounts stained for CD31 and Dll4. Arrowheads point to Dll4^+^ tip cells in the *Vglut1*^-/-^ retina. Scale bars = 41 μm. Students t-test: **<0.02, NS = not significant. Error bars: Mean ± S.E.M.

### Deep plexus vessels are poorly perfused in the *Vglut1*^-/-^ retina

Next, we asked if the deep plexus vessels are polarized and function normally in the *Vglut1*^-/-^ retinas. We stained blood vessels with Podocalyxin, which labels the apical (luminal) side of ECs, and is a marker for endothelial apical-basal polarity **(Figure 3A)** ^34^. Compared to the WT, Podocalyxin coverage of the deep plexus vessels was significantly reduced in *Vglut1*^-/-^ retinas at both P10 and P14 **(Figure 3B, C-E)**, suggesting that EC polarity is abnormal in the deep vascular plexus. Next, we examined whether deep plexus vessels are perfused normally in *Vglut1*^-/-^ retinas. We performed intracardiac injection of fluorescently-labeled Lectin and analyzed its binding to the retinal vasculature as a readout of perfusion. By P14, the entire OPL in *Vglut1*^-/-^ retinas is vascularized **(Figure 1)**. However, the fraction of Lectin^+^ perfused blood vessels was significantly lower in the deep plexus of P14 *Vglut1*^-/-^ compared to the WT retinas **(Figure 3C, F)**. These data demonstrate that, although the deep plexus is fully formed by P14 in the *Vglut1*^-/-^ retinas (**Figure 1F, G**), EC polarity and vessel perfusion remain abnormal.

**Figure 3.**
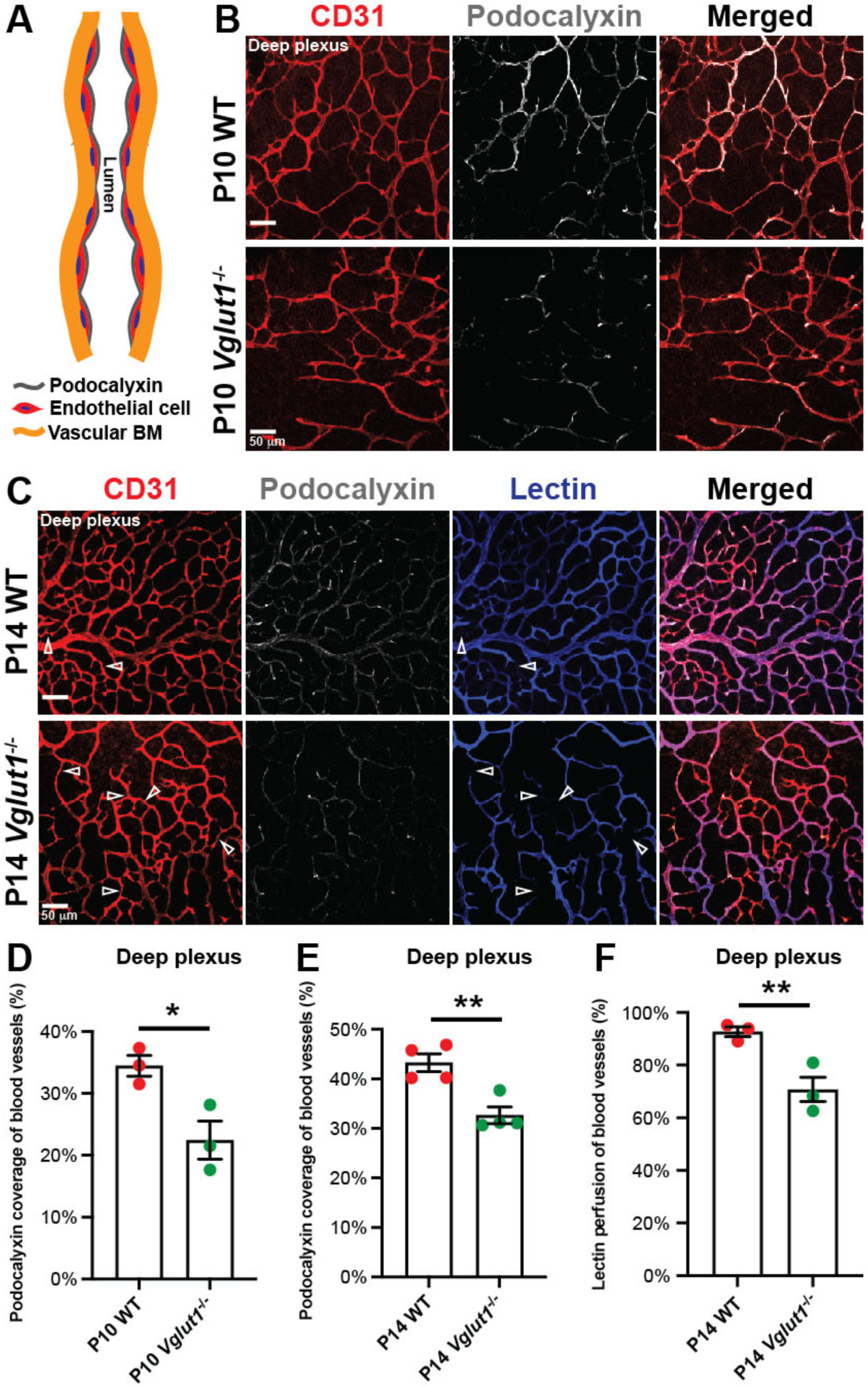
Deep plexus vessels in the *Vglut1*^-/-^ retina are poorly perfused. **A)** Schematic of Podocalyxin localization on the luminal surface of ECs. BM: basement membrane. **B)** Representative images of the deep plexus of P10 WT and *Vglut1*^-/-^ retinal flat mounts labeled with CD31 (red) and Podocalyxin (grey). **C)** Representative images of the deep plexus of P14 WT and *Vglut1*^-/-^ retinal flat mounts labeled with CD31 (red) and Podocalyxin (grey) 5 minutes after intracardiac injection of Lectin (blue). Empty arrowheads show CD31^+^ vessels that are not perfused with Lectin. **D, E)** Dotted bar graphs of Podocalyxin coverage of the deep plexus in P10 (**D**) and P14 (**E**) WT and *Vglut1*^-/-^ retinas (n = 3-4 mice/genotype). **F)** Dotted bar graphs of Lectin perfusion of the deep plexus in P14 WT and *Vglut1*^-/-^ retinas (n=3 mice/genotype). Scale bars = 50 μm. Students t-test: *<0.05, **<0.02, NS = not significant. Error bars: Mean ± S.E.M.

### Paracellular BRB maturation is impaired in the *Vglut1*^-/-^ retina

Next, we asked if diminished glutamate release from photoreceptors and bipolar cells affects BRB maturation. During retinal development, the transcellular (caveolae-mediated) BRB matures by P10 ^35^ whereas the paracellular (tight junction-mediated) BRB does not mature until P18 ^36^ **(Figure 4A)**. We therefore injected intravenously biocytin-tetramethylrhodamine (biocytin-TMR, 870 Da) tracer in WT and *Vglut1*^-/-^ mice at P10 and P14 and quantified its leakage in the retina as a functional readout of paracellular BRB maturation **(Figure 4B)** ^36, 37^. Biocytin-TMR leakage (area and intensity) was higher in the *Vglut1*^-/-^ compared to the WT retinas at both time points **(Figure 4C-I)**. Importantly, these patterns were consistent in both the central and peripheral retina, and the superficial and deep plexuses. Consistent with a delay in BRB maturation, scRNA-seq analyses revealed reduced transcript levels for most “BRB maturation genes” in P10 *Vglut1*^-/-^ compared to WT retinal ECs. These include tight junction-associated genes [*Cldn5* (Claudin-5) and *Tjp1* (ZO-1)], transporters [*Slc7a5* (Lat1), *Slc2a1* (Glut1) etc.,] and effectors of Norrin/β-catenin signaling [*Ctnnb1* and *Lef1*] **(Figure 4J, K; Table S7)**. Surprisingly, transcript levels for *Ocln* (Occludin; tight junction protein), vBM proteins (Collagen IV, Laminin), BBB transcription factors (e.g. *Foxq1*, *Foxf2*, *Zic3* etc.), or caveolar-related transport (*Cav-1*, *Cav-2*, *Cavin-1*, *Mfsd2a*) genes were unaffected in P10 *Vglut1*^-/-^ retinal ECs **(Tables S2, S7)**.

**Figure 4.**
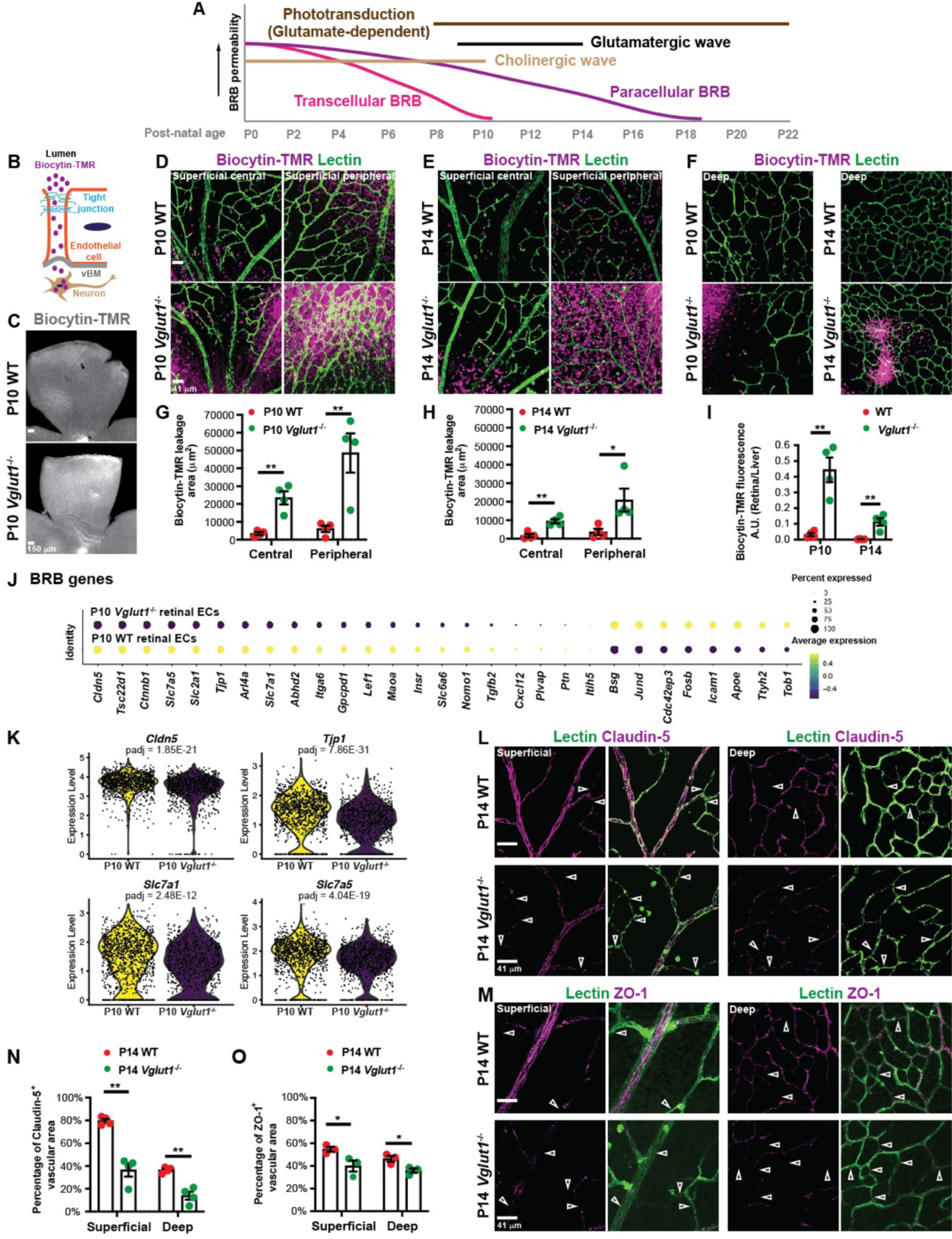
Paracellular BRB maturation is impaired in *Vglut1*^-/-^ retina. **A)** Timeline of the paracellular and transcellular BRB maturation in the postnatal WT retina (P1-P22). **B)** Schematic of biocytin-TMR paracellular leakage through impaired tight junctions at the BRB; the leaked biocytin-TMR is taken up by neurons. vBM = vascular basement membrane. **C)** Representative low magnification images of P10 WT and *Vglut1*^-/-^ retinal flat mounts 40 minutes after intravenous injection of biocytin-TMR, followed by PBS perfusion. **D-F)** Representative high magnification images of the superficial **(D, E)** and deep **(F)** plexuses of P10 and P14 WT and *Vglut1*^-/-^ retinal flat mounts labeled with Lectin (green) 40 minutes after intravenous injection of biocytin-TMR, followed by PBS perfusion. **G-I)** Measurements of the area (**G, H**) and fluorescence intensity (**I**) of leaked biocytin-TMR in P10 and P14 WT and *Vglut1*^-/-^ retinal flat mounts (n = 4). BRB formation is impaired in *Vglut1*^-/-^ retina. **J)** Dot plot of expression levels for BRB genes in P10 WT and *Vglut1*^-/-^ retinal ECs. The dot size indicates the percent of the population expressing each marker; the color scale indicates the average level of gene expression. **K)** Violin plots of representative BRB genes in P10 WT and *Vglut1*^-/-^ retinal ECs. **L)** Representative images of the superficial and deep plexuses of P14 WT and *Vglut1*^-/-^ retinal flat mounts stained for Claudin-5 (Cldn5; purple) and Lectin (green). Empty arrowheads point to Lectin^+^ blood vessels devoid of Claudin-5 at tight junctions. **M)** Representative images of the superficial and deep plexuses in P14 WT and *Vglut1*^-/-^ retinal flat-mounts stained for ZO-1 (*Tjp1*, purple) and Lectin (green). Empty arrowheads point at Lectin^+^ blood vessels devoid of ZO-1 localization to tight junctions. **N, O)** Measurement of Claudin-5 (**N**) and ZO-1 (**O**) coverage of blood vessels in P14 WT and *Vglut1*^-/-^ retinas (n=4 mice / genotype). Scale bars: C = 150 μm; D, L, M = 41 μm. Students t-test: *<0.05, **<0.02. Error bars: Mean ± S.E.M.

We validated the scRNA-seq findings for several tight junction proteins in the *Vglut1*^-/-^ retina. Consistent with the scRNA-seq data, there was a significant reduction in Claudin-5 and ZO-1 protein localization to EC junctions at P14 *Vglut1*^-/-^ compared to the WT retina in both the superficial and deep plexuses by immunofluorescence (**Figure 4L-O**). Importantly, these phenotypes were not caused by abnormalities in the NVU assembly, as mural cell (PDGFRβ^+^), glial endfeet (Aquaporin-4; Aqp-4^+^) and vascular basement membrane (vBM; Laminin) coverage of retinal blood vessels were comparable between the two genotypes **(Figure S2)**.

We next asked whether the lack of glutamate release by neurons affects the maturation of the transcellular BRB properties. We injected intravenously albumin-Alexa594 (MW= 60kDa), a protein that crosses the BRB primarily through a transcellular route ^36^ into P10 WT and *Vglut1*^-/-^ mice and assessed its extravasation into the retina. The tracer was largely confined within the blood vessels and its intensity in the parenchyma was very low and identical between the two genotypes **(Figure S3A, B)**. In addition, there was no difference in endogenous serum IgG leakage between P10 WT and *Vglut1*^-/-^ retinas **(Figure S3C)**. These functional studies are consistent with the scRNA-seq data showing no change in *Mfsd2a* (a gene required for suppression of transcellular BRB permeability ^35^) transcript levels between the two genotypes **(Tables S2, S7)**. In addition, PLVAP expression, a component of the diaphragms of EC fenestrae normally found in the immature BRB and peripheral ECs ^38^, was absent in retinal ECs from both genotypes **(Figure S3D)** despite a slight downregulation in *Plvap* mRNA in P10 *Vglut1*^-/-^ retinal ECs by scRNA-seq **(Figure 4J; Table S7)**. Similarly, immunofluorescence analysis showed no difference in retinal EC expression of Caveolin-1 and Mfsd2a, two critical proteins responsible for regulating EC transcellular permeability ^4, 35, 39, 40^, between the two genotypes **(Figure S3E, F).** Thus, paracellular, but not transcellular, BRB maturation is delayed in both the superficial and deep plexuses of *Vglut1*^-/-^ retinas with decreased extracellular glutamate **(Figure S3G)**.

### BRB maturation is accelerated in the absence of cholinergic waves due to an early onset of glutamatergic activities

During development, the cholinergic waves persist beyond P10 in the *Vglut1*^-/-^ retina lacking glutamatergic neuronal activities ^12^. Conversely, the glutamatergic waves begin earlier (P8) than usual (P10) in the *Chrnb2*^-/-^ retina where the cholinergic waves are abolished (**Figure 5A**) ^11^. Recently it was reported that pharmacological blockade of cholinergic waves before P8 delays deep plexus angiogenesis and BRB maturation, suggesting a critical role for the cholinergic waves in promoting these processes in the developing retina ^14^. In contrast, our phenotypic analysis of the *Vglut1*^-/-^ retinal vasculature implies that the cholinergic waves are not sufficient to induce paracellular BRB maturation since it is very delayed in the *Vglut1*^-/-^ retina throughout P14 (**Figure 4**). To test this hypothesis, we examined angiogenesis and the paracellular BRB integrity in the *Chrnb2*^-/-^retina where cholinergic waves are abolished and glutamatergic waves begin earlier (P8) than usual ^11^ **(Figure 5A)**. At P7, before the onset of glutamatergic waves in the *Chrnb2*^-/-^ retina, there was no difference in the vascularized area of the superficial plexus **(Figure 5B, C),** and biocytin-TMR leakage **(Figure 5D-F)** between WT and *Chrnb2*^-/-^ retinas. In contrast, there was less biocytin-TMR leakage in the parenchyma **(Figure 5J-L),** and increased Claudin-5 coverage of blood vessels **(Figure 5M, N)** in the P10 *Chrnb2*^-/-^ retina compared to the WT. However, vascular coverage in either the superficial or deep plexus was not significantly different between P10 WT and *Chrnb2*^-/-^ retinas **(Figure 5G-I)**. These data are consistent with the idea that the earlier onset of glutamatergic waves in the *Chrnb2*^-/-^ retina is sufficient to promote a precocious BRB maturation, but does not affect deep plexus angiogenesis. Moreover, they demonstrate that glutamatergic neuronal activities, but not the cholinergic waves, are the primary drivers of paracellular BRB maturation in the developing retina.

**Figure 5.**
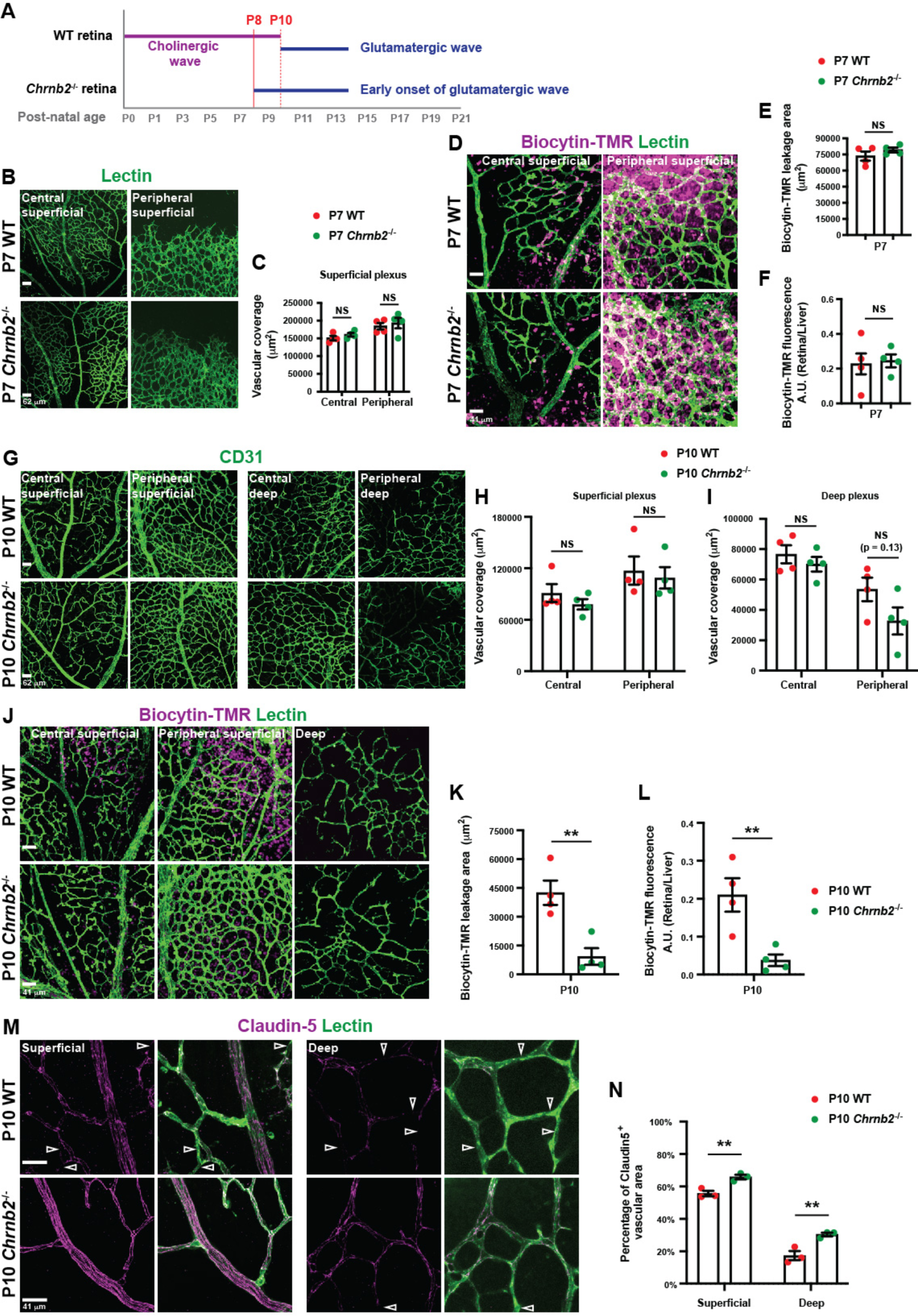
Paracellular BRB matures precociously in the *Chrnb2*^-/-^ retina. **A)** Schematic diagram illustrating that glutamatergic wave begins earlier (P8) than usual (P10) in the *Chrnb2*^-/-^ retina where cholinergic wave is abolished. **B)** Representative images of the superficial plexus of P7 WT and *Chrnb2*^-/-^ retinal flat-mounts labeled with Lectin (green). **C)** Quantification of vascular coverage in the superficial plexus of P7 WT and *Chrnb2*^-/-^ retinas. **D)** Representative images of superficial and deep plexuses of P7 WT and *Chrnb2*^-/-^ retinal flat-mounts labeled with Lectin (green) 40 minutes after intravenous biocytin-TMR injection, followed by PBS perfusion. **E, F)** Measurements of the area (**E**) and fluorescence intensity (**F**) of leaked biocytin-TMR in P7 WT and *Chrnb2*^-/-^ retinal flat-mounts (n = 4 mice / genotype). **G)** Representative images of superficial and deep plexuses of P10 WT and *Chrnb2*^-/-^ retinal flat-mounts labeled with CD31 (green). **H, I)** Quantification of vascular coverage in the superficial (**H**) and deep (**I**) plexuses of P10 WT and *Chrnb2*^-/-^ retinas. **J)** Representative images of superficial and deep plexuses of P10 WT and *Chrnb2*^-/-^ retinal flat-mounts labeled with Lectin (green) 40 minutes after intravenous biocytin-TMR injection, followed by PBS perfusion. **K, L)** Measurements of the area (**K**) and fluorescence intensity (**L**) of leaked biocytin-TMR in P10 WT and *Chrnb2*^-/-^ retinal flat-mounts (n = 4 mice / genotype). **M)** Representative images of the superficial and deep plexuses of P10 WT and *Chrnb2*^-/-^ retinal flat-mounts stained for Claudin-5 (purple) and Lectin (green). Empty arrowheads point at Lectin^+^ blood vessels devoid of Claudin-5 localization to tight junctions. **N)** Measurement of Claudin-5 coverage of blood vessels in P10 WT and *Chrnb2*^-/-^ retinas (n = 3 mice / genotype). Scale bars: B, G = 62 μm, D, J, M = 41 μm. Students t-test: **<0.02. Error bars: Mean ± S.E.M.

### Deep plexus angiogenesis and paracellular BRB maturation are accelerated in the *Gnat1*^-/-^ retina

To further evaluate the effect of excess glutamate release by retinal neurons, we examined the retinal vascular phenotypes in mice lacking α−Transducin. Phototransduction by rod photoreceptors is a complex cascade, dependent in part on the Gα subunit, Transducin 1 (*Gnat1*). *Gnat1*^-/-^ rod photoreceptors remain constitutively depolarized ^26, 27^ leading to increased glutamate release from rods **(Figure 6A, B)**. To understand how excess glutamate release by rod photoreceptors affects retinal angiogenesis, we compared the vascular growth of the superficial and deep plexuses between the WT and *Gnat1*^-/-^ retinas. While the vascular coverage of the retina in the superficial plexus was similar between the two genotypes at P10 **(Figure 6C, D)**, both the vascular coverage of the retina in the deep plexus, and its branch-point density were significantly increased in the *Gnat1*^-/-^ retina **(Figure 6E-J)**. However, the angiogenesis phenotype was transient, and by P14 there was no difference between the two genotypes **(Figure 6F, G, I, J; Figure S4A)**. Importantly, these patterns were similar in both central and peripheral retina. Consistent with an accelerated deep plexus angiogenesis in the P10 *Gnat1*^-/-^, there was also increased EC proliferation (Ki67^+^ ECs) in mutant deep vascular plexus compared to the WT **(Figure 6K, L)**.

**Figure 6.**
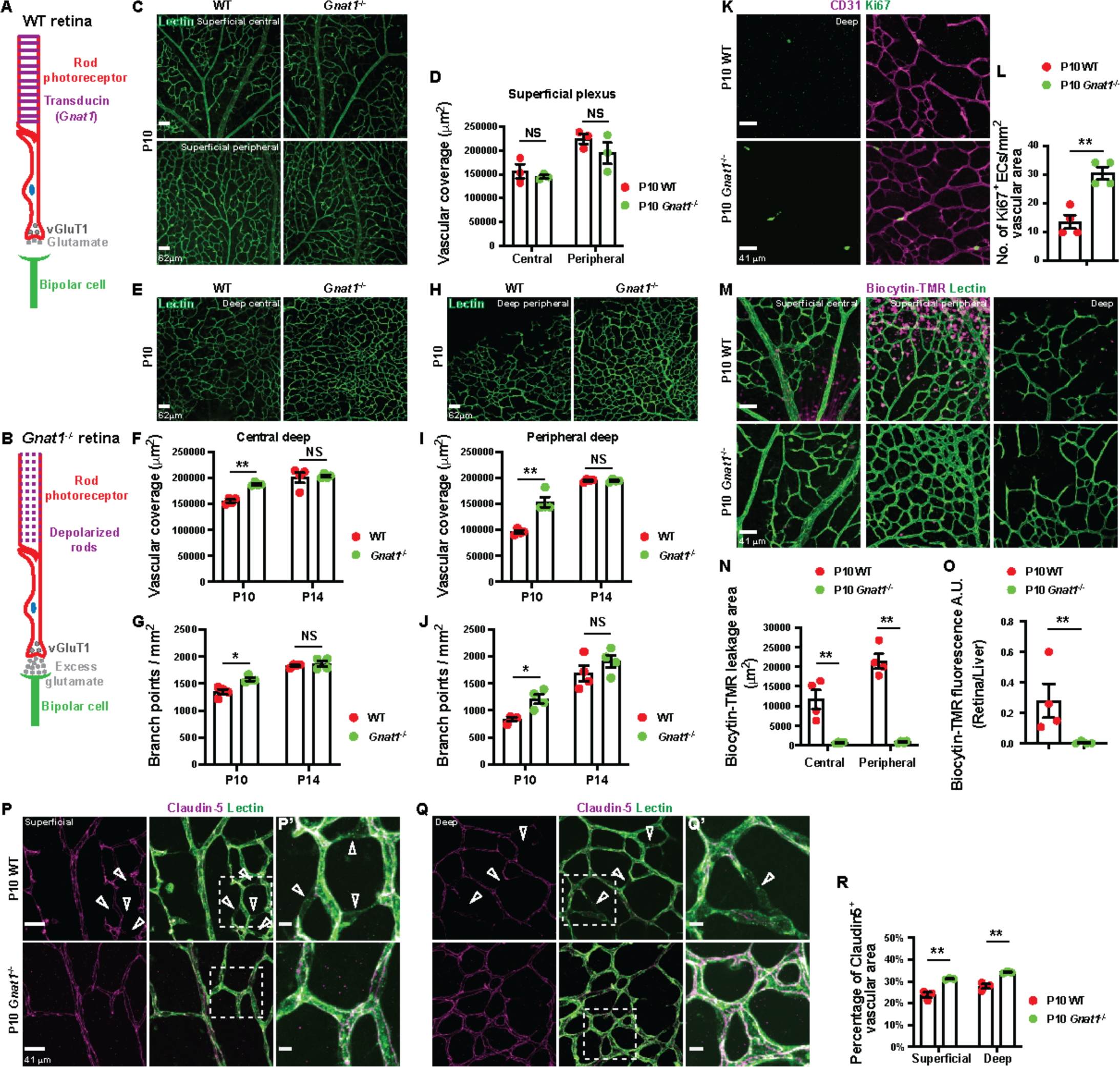
Deep plexus angiogenesis and paracellular BRB maturation are accelerated in the *Gnat1*^-/-^ retina. **A, B)** Schematic diagrams of glutamate release from rod photoreceptors in WT (**A**) and *Gnat1*^-/-^ (**B**) retinas. *Gnat1*^-/-^ rod photoreceptors (depolarized) release excess glutamate into the synaptic cleft. **C)** Representative images of the superficial plexus in P10 WT and *Gnat1*^-/-^ retinal flat-mounts stained for Lectin (green). **D)** Measurements of vascular coverage of the superficial plexus in P10 WT and *Vglut1*^-/-^ central and peripheral retina (n = 3 mice / genotype). **E, H)** Representative images of the deep plexus in P10 WT and *Gnat1*^-/-^ retinal flat-mounts stained for Lectin (green). **F, G, I, J)** Measurements of vascular coverage (**F, I**) and branch point density (**G, J**) of the deep plexus in P10 and P14 WT and *Gnat1*^-/-^ central (**F, G**) and peripheral (**I, J**) retina (n = 4 mice / genotype). **K)** Representative images of the deep plexus in P10 WT and *Gnat1*^-/-^ retinal flat-mounts stained for Ki67 (purple) and CD31 (green). **L)** Measurements of Ki67^+^ ECs in P10 WT and *Gnat1*^-/-^ retinal deep plexus (n = 4 mice / genotype). **M)** Representative images of superficial and deep plexuses of P10 WT and *Gnat1*^-/-^ retinal flat-mounts labeled with Lectin (green) 40 minutes after intravenous injection of biocytin-TMR (purple), followed by PBS perfusion. **N, O)** Measurements of the area (**N**) and fluorescence intensity (**O**) of leaked biocytin-TMR in P10 WT and *Gnat1*^-/-^ retinal flat-mounts (n = 4 mice / genotype). **P, Q)** Representative images of the superficial (**P**) and deep (**Q**) plexuses in P10 WT and *Gnat1*^-/-^ retinal flat-mounts stained for Claudin-5 (purple) and Lectin (green). P’ and Q’ are magnified images of the corresponding boxed areas. Empty arrowheads point at Lectin^+^ blood vessels devoid of Claudin-5 localization to tight junctions. **R)** Measurement of Claudin-5 coverage of blood vessels in P10 WT and *Gnat1*^-/-^ retinas (n = 4 mice / genotype). Scale bars: **C, E, H** = 62 μm and **K, M, P, Q** = 41 μm. Students t-test: *<0.05, **<0.02, NS = not significant. Error bars: Mean ± S.E.M.

Next, we asked whether excess glutamate release by rod photoreceptors affects paracellular BRB maturation. Biocytin-TMR leakage area and fluorescence intensity were reduced in *Gnat1*^-/-^ compared to the WT retinas in both central and peripheral retinas and superficial and deep plexuses at P10 and P14 **(Figures 6M-O; S4B-D)**. Consistent with the functional data, there was a significant increase in Claudin-5 protein localization at BRB tight junctions in the P10 *Gnat1*^-/-^ retina compared to the WT **(Figure 6P-R)**. In summary, retinal deep plexus angiogenesis is accelerated and paracellular BRB integrity is precociously mature when there is excess glutamate release by the rods.

To address whether aberrant specification of neuronal subtypes or Müller glia in either *Vglut1*^-/-^ or *Gnat1*^-/-^ retinas may be responsible for the vascular phenotypes seen in these mutants, we immunolabelled and quantified the number of neuronal subtypes and Müller glia. There was no significant difference in the number of ganglion cells [NeuN^+^ ^41, 42^], amacrine (except for a slight reduction in the *Vglut1*^-/-^ retina) and horizontal cells [Calbindin^+^ ^43^], photoreceptors [measured by outer nuclear layer (ONL) thickness] among the three genotypes **(Figure S5A-I)**. Similarly, the number of Sox9^+^ Müller glia ^44, 45^ was similar among WT, *Vglut1*^-/-^ and *Gnat1*^-/-^ retinas **(Figure S5J, K)**. Importantly, altered levels of extracellular glutamate did not trigger Müller cell gliosis in *Vglut1*^-/-^ and *Gnat1*^-/-^ retinas, as there was no upregulation of GFAP expression [a marker of gliosis ^46, 47^] in Müller glia processes spanning multiple retinal layers (**Figure S5L**). GFAP expression was only detected in retinal astrocytes over the GCL (colocalized with PDGFRα) **(Figure S5L, M)**. Therefore, the angiogenesis and BRB phenotypes seen in *Vglut1*^-/-^ and *Gnat1*^-/-^ retinas are not caused by defects in neuronal and glial cell type specification.

Finally, we examined whether cholinergic, GABAergic and dopaminergic neuronal populations are altered in mutant retinas with abnormal glutamatergic activities which may indirectly affect vascular development. There was no difference in the number of ChAT^+^ (cholinergic), GAD67^+^ (GABAergic) and Tyrosine Hydroxylase (TH)^+^ (dopaminergic) neurons between WT, *Vglut1*^-/-^ and *Gnat1*^-/-^ retinas **(Figure S6A, B)**. Overall, our data show that changes in extracellular glutamate does not affect neuronal or glial subtypes specification or neurons involved in other types of neurotransmission.

### Neurons and Müller glia in the inner nuclear layer of the retina respond to altered levels of extracellular glutamate

To understand how extracellular glutamate affects retinal angiogenesis and paracellular BRB maturation, we examined several parameters that could influence these processes. First, we asked whether glutamate acts directly on retinal ECs. Our scRNA-seq analyses of P10 and P14 WT retinal ECs showed that neither ionotropic (*Grin1*, *Grin2b*, *Grin2d*, *Gria1*), nor metabotropic (*Grm3*, *Grm5*, *Grm6*, *Grm7*, *Grm8*) glutamate receptor subunit transcripts are expressed in developing retinal ECs **(Figure S6C).** Analysis of a previously published scRNA-seq database of all cell types from the P14 retina ^48^ also confirmed our findings that glutamate receptor subunit transcripts are not expressed in P14 retinal ECs. Thus, glutamate cannot act directly on developing retinal ECs. In contrast, ionotropic and metabotropic glutamate receptor subunit mRNAs are expressed by the inner nuclear layer (INL) neurons and Müller glia **(Figure S6D)**, suggesting that extracellular glutamate could act on INL neurons and Müller glia.

To examine whether altered levels of extracellular glutamate are sensed by Müller glia in *Vglut1*^-/-^ and *Gnat1*^-/-^ retinas, we initially measured mean fluorescence intensity of glutamine synthetase (GS), an enzyme that coverts glutamate to glutamine, and the Glutamate Aspartate Transporter [GLAST, excitatory aminoacid transporter 1 (EAAT-1)] from immunofluorescence staining. These proteins are known to be upregulated in Müller glia exposed to excess extracellular glutamate ^49, 50^. GS and GLAST protein levels were downregulated in *Vglut1*^-/-^, and upregulated in *Gnat1*^-/-^ retinas, compared to the WT retinas **(Figure 7A-D)**. To determine whether altered levels of extracellular glutamate are sensed by neurons in *Vglut1*^-/-^ and *Gnat1*^-/-^ retinas, we measured Glutamate Transporter 1 protein levels [GLT-1, or excitatory aminoacid transporter 1 (EAAT-2)]. This transporter is expressed primarily in retinal neurons ^51^, and its expression is regulated by extracellular glutamate ^52^. GLT-1 expression in INL neurons was downregulated in the *Vglut1*^-/-^, and upregulated in the *Gnat1*^-/-^ retinas, compared to the WT **(Figure 7E-F)**. These data confirm that Müller glia and INL neurons respond to abnormal extracellular glutamate levels in *Vglut1*^-/-^ or *Gnat1*^-/-^ retinas.

**Figure 7.**
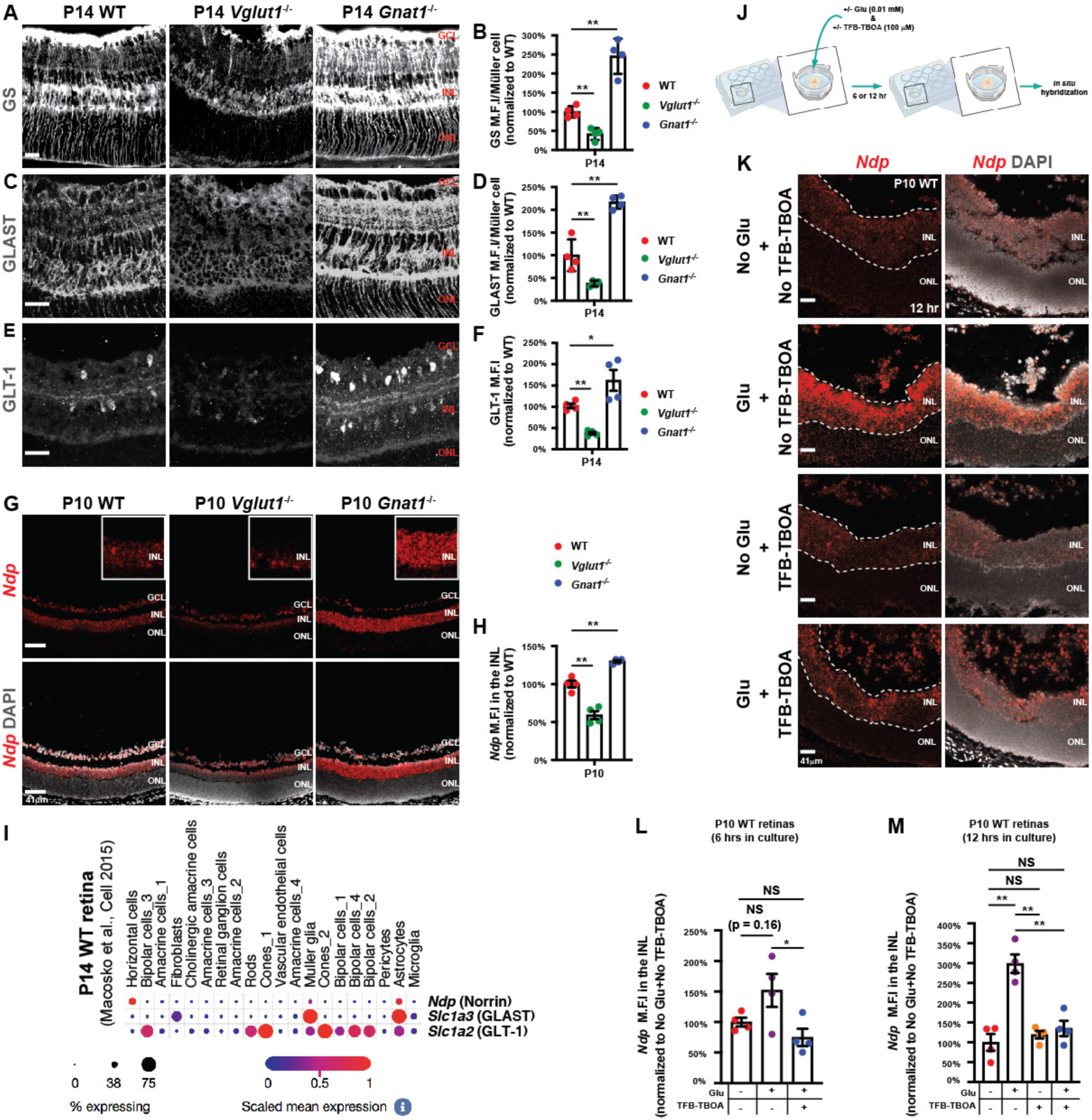
Expression of glutamine synthetase, glutamate transporters and Norrin are downregulated in *Vglut1*^-/-^ retinas and upregulated in *Gnat1*^-/-^ retinas. **A, B)** P14 WT, *Vglut1*^-/-^ and *Gnat1*^-/-^ retinal sections were stained for glutamine synthetase (GS) (**A**), and the mean fluorescence intensity (M.F.I) in the INL was quantified (**B**) for each genotype (n = 4 mice / genotype). **C, D)** P14 WT, *Vglut1*^-/-^ and *Gnat1*^-/-^ retinal sections were stained for GLAST (**C**), and the M.F.I in the INL was quantified (**D**) for each genotype (n = 4 mice / genotype). **E, F)** P14 WT, *Vglut1*^-/-^ and *Gnat1*^-/-^ retinal sections were stained for GLT-1 (**E**), and the M.F.I in the INL was quantified (**F**) for each genotype (n=4 mice / genotype). **G)** Fluorescent *in situ* hybridization of P10 WT, *Vglut1*^-/-^ and *Gnat1*^-/-^ retinal sections with an antisense probe against *Ndp* (Norrin), followed by staining with DAPI. Norrin is expressed throughout the INL. **H)** Quantification of the M.F.I of *Ndp* in the INL of P10 WT, *Vglut1*^-/-^ and *Gnat1*^-/-^ retinal sections (n = 4 mice / genotype). **I)** Expression profiles of *Ndp*, *Slc1a3* (GLAST) and *Slc1a2* (GLT-1) in the P14 WT retina from an existing scRNA-seq database ^48^. *Ndp* mRNA is expressed predominantly by the horizontal cells and Müller glia in the INL (small subsets of amacrine and bipolar cells also express *Ndp*). In the INL, *Slc1a3* is primarily expressed by Müller glia and astrocytes, whereas *Slc1a2* is expressed in retinal neurons as well as Müller glia. **J)** Schematic of *ex vivo* organotypic culture experiments. **K)** Fluorescent *in situ* hybridization of P10 WT retinal sections, cultured *ex vivo* for 12 hours with or without glutamate or TFB-TBOA in the media, with an antisense probe against *Ndp* (*Norrin*) mRNA, followed by staining with DAPI. *Norrin* mRNA is expressed throughout the INL. **L, M)** Quantification of the M.F.I of *Ndp* mRNA in the INL of P10 WT retinal sections, cultured *ex vivo* for either 6 **(L)** or 12 **(M)** hours with or without glutamate or TFB-TBOA in the media (n = 4 retinas / group). GCL: ganglion cell layer, INL: inner nuclear layer, ONL: outer nuclear layer. Scale bars = 41 μm. One-way ANOVA: *<0.05, **<0.02, NS = not significant. Error bars: Mean ± S.E.M.

### Retinal metabolic demand, hypoxia and VEGF-A signaling are not affected in the *Vglut1*^-/-^ retina

Since Müller glia and INL neurons respond to changes in extracellular glutamate levels, we asked whether neuronal metabolic demand is also affected in *Vglut1*^-/-^ and *Gnat1*^-/-^ retinas, as alterations in neuronal metabolic demand trigger vascular abnormalities [reviewed in ^4^]. We examined Hexokinase-2 (HK2) and Pyruvate Kinase M-2 (PKM2) protein levels by Western blotting since they are key glycolytic enzymes of neuronal metabolism readout ^53^. The levels of these enzymes were similar among P10 WT, *Gnat1*^-/-^ and *Vglut1*^-/-^ retinas **(Figure S7A-L)**. Thus, alterations in extracellular glutamate levels caused by genetic perturbations in glutamate release from neurons do not affect expression of key enzymes for the neuronal metabolism in the retina.

Neuronal-derived VEGF-A is critical for nervous system function ^54^ and VEGF-A signaling is a major regulator of retinal angiogenesis ^14, 55^, prompting us to examine whether VEGF-A levels change in *Vglut1*^-/-^ and *Gnat1*^-/-^ retinas. VEGF-A protein levels measured with Western blotting were not different among WT, *Vglut1*^-/-^ and *Gnat1*^-/-^ retinas **(Figure S7M-P)**. These results are also consistent with our scRNA-seq data showing no changes in transcripts for VEGF-A signaling components [*Nr4a2*, *Egr3*, *Igfbp3*, *Hlx1*, *Crem*, *Per1*, *Mef2c*, *Thbd*, *Ndrg1*, *Dnajb9*, *Mycn* ^56^] in *Vglut1*^-/-^ retinal ECs **(Figure S7Q; Table S2)**. Since VEGF-A expression depends on tissue hypoxia ^57^, we measured retinal hypoxia with a hypoxyprobe, Pimonidazole ^58, 59^. Consistent with no change in VEGF-A protein and downstream target transcript levels, there was no difference in tissue hypoxia between *Vglut1*^-/-^ and WT retinas **(Figure S7R, S)**. Therefore, the vascular phenotypes observed in mutants with altered extracellular glutamate are not due to changes in VEGF-A signaling and hypoxia.

### Extracellular glutamate levels affect Norrin expression in retinal neurons and Müller glia and as a consequence endothelial Norrin/β-catenin signaling

Our scRNA-seq analyses showed that transcript levels for two Norrin/β-catenin signaling effectors (*Ctnnb1* and *Lef1*), essential for angiogenesis and BRB maturation, were downregulated in P10 *Vglut1*^-/-^ compared to WT retinal ECs **(Figures 2D, F; 4J)**. Among seven key transcription factors (*Foxf2*, *Foxl2*, *Foxq1*, *Lef1*, *Ppard*, *Zfp551* and *Zic3*) required for BBB maturation ^60^, only *Lef1* was downregulated significantly in P10 *Vglut1*^-/-^ retinal ECs, suggesting a specific effect on Norrin/β-catenin signaling. To address changes in Norrin/β-catenin signaling, we examined *Norrin* (*Ndp*) mRNA expression in *Vglut1*^-/-^ and *Gnat1*^-/-^ retinas by fluorescent *in situ* hybridization (FISH). *Norrin* mRNA expression was reduced in the INL of P10 *Vglut1*^-/-^ retinas, and upregulated in the INL of P10 *Gnat1*^-/-^ retinas, compared to the WT **(Figure 7G, H)**. The upregulation of *Norrin* mRNA persisted in the INL of *Gnat1*^-/-^ retinas through P14 **(Figure S4E)**. Consistent with our data, analysis of a previously published scRNA-seq database ^48^ demonstrated that INL neurons (horizontal cells and a subset of amacrine and bipolar cells) and Müller glia express high levels of *Norrin* mRNA and glutamate transporters [*Slc1a2* (GLT-1) and *Slc1a3* (GLAST)] respectively ^48^ **(Figure 7I)**. These transcriptional data and the response of the INL neurons and Müller glia to extracellular glutamate in the *Vglut1*^-/-^ and *Gnat1*^-/-^ retinas (**Figure 7A-F**) suggest that glutamate may act directly on Müller glia and INL neurons to induce *Norrin* mRNA expression.

To test whether glutamate can induce *Norrin* mRNA expression in Müller glia and INL neurons, we performed *ex vivo* organotypic cultures of P10 WT retinas in the presence or absence of glutamate (0.01mM) and measured *Norrin* mRNA expression in the INL by FISH (fluorescence intensity) after 6 and 12 hours in culture (**Figure 7J)**. *Norrin* mRNA expression in the INL was upregulated after 6 hours of glutamate exposure, although it was not statistically significant (p=0.16; **Figure 7K, L)**. By 12 hours, *Norrin* mRNA levels in the INL were upregulated significantly with glutamate compared to media alone **(Figure 7K, M)**. Therefore, glutamate induces *Norrin* mRNA expression in Müller glia and INL neurons.

Glutamate-mediated induction of *Norrin* mRNA expression in Müller glia and INL neurons may be regulated through glutamate uptake via GLAST and GLT-1. To test this directly, we performed *ex vivo* organotypic cultures of P10 WT retinas with or without glutamate and TFB-TBOA, a pharmacological inhibitor of GLAST and GLT-1^61–63^ **(Figure 7J)**. *Norrin* mRNA expression in the INL was downregulated significantly with glutamate and TFB-TBOA compared to glutamate alone conditions after 6 and 12 hours in culture **(Figure 7K-M)**. Importantly, TFB-TBOA alone did not alter *Norrin* mRNA levels in the INL compared to the media alone **(Figure 7K, M)**, suggesting that TFB-TBOA did not have any non-specific effect on *Norrin* mRNA expression. Based on these pharmacological data, we conclude that glutamate uptake by GLAST and GLT-1 likely induces *Norrin* mRNA expression in both Müller glia and INL neurons.

Next, we examined Norrin/β-catenin signaling activation in *Vglut1*^-/-^ and *Gnat1*^-/-^ retinal ECs. We immunolabeled P14 WT, *Vglut1*^-/-^ and *Gnat1*^-/-^ retinal sections with Lef1 and quantified the number of Lef1^+^ ECs in both the superficial and deep plexuses. There were significantly fewer Lef1^+^ ECs in the *Vglut1*^-/-^ retina and significantly higher in the *Gnat1*^-/-^ retina compared to the WT **(Figure 8A, B)**. These data suggest that angiogenesis and BRB phenotypes in *Vglut1*^-/-^ and *Gnat1*^-/-^ retinas are likely driven by altered Norrin/β-catenin signaling.

**Figure 8.**
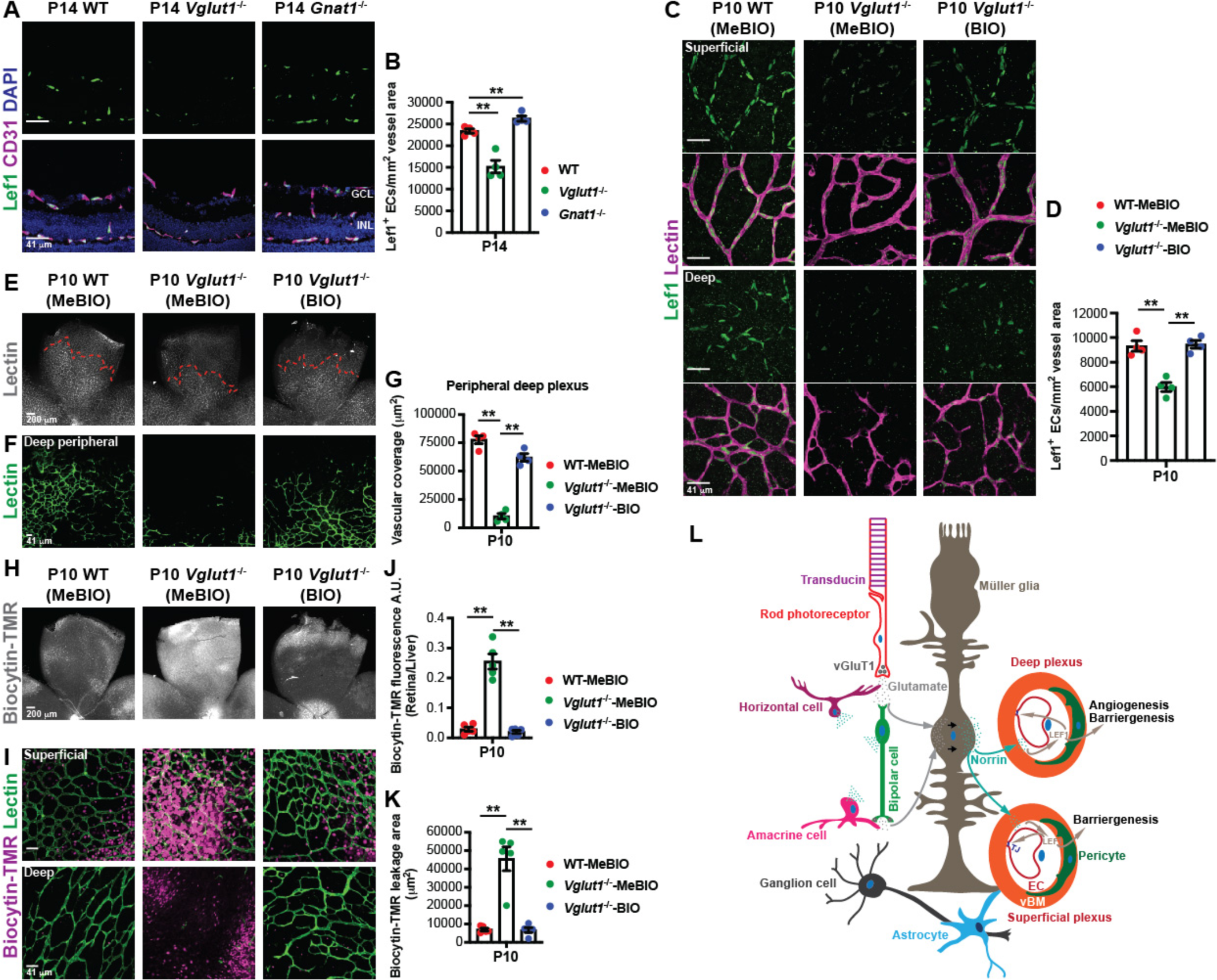
Upregulation of Norrin/β-catenin signaling rescues vascular deficits of the *Vglut1*^-/-^ retina. **A, B)** P14 WT, *Vglut1*^-/-^ and *Gnat1*^-/-^ retinal sections were stained (**A**) for Lef1 (green), CD31 (purple) and DAPI (blue), and the number of Lef1^+^ ECs was quantified (**B**) for each genotype (n = 4 mice / genotype). **C, D)** MeBIO (control)- or BIO (activates Norrin/β-catenin signaling)-treated P10 WT and *Vglut1*^-/-^ retinal flat-mounts were stained (**C**) with Lectin (purple) and Lef1 (green), and the number of Lef1^+^ ECs / vessel area (mm^2^) for each treatment was quantified (**D**) for each treatment (n = 4 mice / group). BIO activates Norrin/β-catenin signaling in ECs of *Vglut1*^-/-^ retinas. **E-G)** MeBIO- or BIO-treated P10 WT and *Vglut1*^-/-^ retinas were stained with Lectin [low (grey; **E**) and high (green; **F**) magnifications], and vascular coverage of the peripheral deep plexus was quantified (**G**) for each treatment (n = 4 mice / group). Red dotted lines depict the extent of deep plexus growth. **H, I)** Representative images of flat-mount MeBIO- or BIO-treated P10 WT and *Vglut1*^-/-^ retinas 40 minutes after intravenous injection of biocytin-TMR [low (grey; **H**) and high (purple; **I**) magnifications] and stained with Lectin (green). P10 *Vglut1*^-/-^ retinas treated with MeBIO have high biocytin-TMR leakage. P10 *Vglut1*^-/-^ retinas treated with BIO have low biocytin-TMR leakage similar to P10 WT retinas. **J, K**) Dotted bar graphs show quantification of biocytin-TMR fluorescence intensity (**J**) and leakage area (**K**) (n = 5 mice / group). **L)** According to our working model, glutamate is taken up into synaptic vesicles by vGluT1 (*Vglut1*), and is released into the synaptic cleft by neurons, either during spontaneous glutamatergic waves (from bipolar cells) or light-dependent glutamatergic synaptic activities of photoreceptors in the retina. The inhibition of glutamate release from rods after light exposure is regulated by Transducin (*Gnat1*). The released glutamate signals to Müller glia and inner nuclear layer (INL) neurons including horizontal, amacrine and bipolar cells. As a result, Müller glia and INL neurons upregulate Norrin (*Ndp*) expression and likely its secretion; however, the relative contribution from Müller glia and INL neurons on Norrin levels in this process is unclear. Glutamate-dependent secretion of Norrin activates endothelial Norrin/β-catenin signaling, which in turn promotes deep plexus angiogenesis and the paracellular BRB maturation in both the superficial and deep plexuses of the retina. GCL: ganglion cell layer, INL: inner nuclear layer. Scale bars **A, D, G** = 41 μm, **C, F** = 200 μm. One-way ANOVA: **<0.02. Error bars: Mean ± S.E.M.

### Activation of endothelial Norrin/β-catenin signaling restores retinal angiogenesis and BRB deficits in *Vglut1^-/^*^-^ mice

If angiogenesis and BRB phenotypes observed in *Vglut1*^-/-^ and *Gnat1*^-/-^ retinas are driven by altered endothelial Norrin/β-catenin activity, then activation of Norrin/β-catenin signaling in retinal ECs should rescue angiogenesis and BRB phenotypes in *Vglut1*^-/-^ retinas. We injected intraperitoneally either BIO, a compound that activates Wnt (Norrin)/β-catenin signaling by inhibiting GSK-3β, or an inactive analog MeBIO ^64^, in WT and *Vglut1*^-/-^ mice between P6 - P8 and analyzed the vascular and BRB phenotypes in the retina at P10. BIO-treatment significantly increased the number of Lef1^+^ ECs in the *Vglut1*^-/-^ retina compared to the MeBIO-treated *Vglut1*^-/-^ retina, whereas MeBIO administration had no effect on endothelial Lef1 [there were significantly fewer Lef1^+^ ECs in the MeBIO-treated *Vglut1*^-/-^ compared to the MeBIO-treated WT retina (**Figure 8C, D**)]. Importantly, there were no Lef1^+^ nuclei outside the vasculature with either treatment **(Figure 8C, D)**. Therefore, BIO-treatment activated Norrin/β-catenin signaling specifically in *Vglut1*^-/-^ retinal ECs.

BIO-treatment significantly increased vascular coverage by the deep plexus in *Vglut1*^-/-^ retina to levels comparable to MeBIO-treated WT retinas **(Figure 8E-G)**. However, the vascular coverage of the deep plexus was significantly lower in the MeBIO-treated *Vglut1*^-/-^ retina compared to the MeBIO-treated WT retina, consistent with no effect of MeBIO on endothelial Norrin/β-catenin activation (**Figure 8C, D**). Similarly, BIO-treatment significantly decreased biocytin-TMR leakage area and intensity in the *Vglut1*^-/-^ retina similar to those observed in MeBIO-treated WT levels **(Figure 8H-K),** whereas Me-BIO treatment had no effect [biocytin-TMR leakage area and intensity were significantly higher in the MeBIO-treated *Vglut1*^-/-^ retina compared to the MeBIO-treated WT retina (**Figure 8H-K**)]. Interestingly, although BIO administration in WT mice decreased modestly biocytin-TMR leakage in the retina compared to the MeBIO administration **(Figure S8C-E)**, the vascular coverage of the deep plexus was reduced significantly in the BIO-treated compared to the MeBIO-treated WT retina **(Figure S8A, B)**. These data are consistent with our previous findings that endothelial Norrin/β-catenin activation needs to be finely tuned for proper deep plexus angiogenesis and BRB maturation in the developing retina ^36^. In summary, activation of endothelial Norrin/β-catenin signaling was sufficient to rescue angiogenesis and BRB phenotypes in the *Vglut1*^-/-^ retina.

Taken together, our data demonstrate that glutamate released by neurons regulates retinal angiogenesis and paracellular BRB integrity by modulating Norrin expression in retinal neurons and Müller glia and endothelial Norrin/β-catenin signaling activation.

## DISCUSSION

It is well established that neuronal and vascular development in the CNS, including the retina, are interdependent. Axonal growth and sprouting angiogenesis use multiple similar guidance cues ^65, 66^. CNS vascular growth is also regulated by the neuronal metabolism ^43, 67, 68^, as CNS is the most metabolically active organ system in the body. CNS neurons use a multitude of neurotransmitters such as glutamate, acetyl choline, dopamine and GABA. However, it is unclear whether specific types of neuronal activities and neurotransmitters affect CNS vascular growth. A recent study suggested that dopaminergic activities of ganglion cells regulate retinal superficial plexus angiogenesis ^69^. Cholinergic waves propagated by starburst amacrine cells have also been implicated in the development of retinal deep plexus vessels and BRB maturation ^14^. While glutamate is the predominant neurotransmitter in the retina during vascular development and in adulthood ^70^, nothing is known about whether glutamatergic neuronal activities regulate retinal angiogenesis and BRB maturation. Our study elucidates this critical missing step that links glutamatergic neuronal activities with vascular development and maturation in the retina - a process that may also be relevant for several retinal disease. Below, we discuss: a) the role of glutamate in angiogenesis and barriergenesis, b) the signaling pathways that function downstream of glutamate to promote these processes, and c) the relevance for the disease.

In our study, we used three mouse models to address the role of glutamate in angiogenesis and barriergenesis: a) *Vglut1*^-/-^, where retinal photoreceptors and bipolar cells fail to release glutamate into the synaptic cleft, b) *Gnat1*^-/-^, where depolarized rod photoreceptors release excess glutamate into the synaptic cleft, and c) *Chrnb*2^-/-^, where cholinergic waves are abolished but glutamatergic waves begin at P8 instead of P10. We found that deep plexus angiogenesis and paracellular BRB maturation in the retina are critically linked to the extracellular glutamate levels. These processes are delayed in *Vglut1*^-/-^ retinas, where neurons fail to release glutamate. In contrast, deep plexus angiogenesis and paracellular BRB maturation are accelerated in *Gnat1*^-/-^ retinas, where constitutively depolarized rods release excessive glutamate. Although the deep plexus angiogenesis, which occurs from P8-P12, was affected in all mutant mouse models, the superficial plexus angiogenesis (P0-P8) was unperturbed. This is likely because the superficial plexus is fully formed by P8, before the onset of glutamatergic neuronal activities in the mouse retina. This developmental time course can also explain why treatment with 2-amino-4-phosphonobutyrate, a compound that blocks glutamatergic waves, from P3-P7 did not affect retinal angiogenesis in a previous study ^14^, because the treatment was performed before the onset of glutamatergic waves in the developing mouse retina. Similarly, we found that the transcellular, but not paracellular BRB, was also not affected in the *Vglut1*^-/-^ retina. This is consistent with previous studies showing that transcellular BRB in the mouse retina is fully mature by P10 ^3^, whereas paracellular BRB does not fully mature until P18 ^36^. Thus, it is conceivable that glutamatergic neuronal activities kick in too late to affect the maturation of the transcellular BRB, but have enough time to influence the maturation of the paracellular BRB.

Interestingly, *Vglut1*^-/-^ retinas lacking glutamatergic neuronal activities show persistent cholinergic waves beyond P10 ^12^. Since deficits in the deep plexus angiogenesis and paracellular BRB integrity persist in the *Vglut1*^-/-^ retina throughout P14, our data suggest that the cholinergic waves are not sufficient to promote deep plexus angiogenesis and paracellular BRB maturation. In contrast, paracellular BRB is precociously mature in the *Chrnb2*^-/-^ retina where cholinergic waves are abolished and glutamatergic waves begin earlier (P8) than usual ^11^. Overall these data indicate that glutamatergic neuronal activities are the primary driver of paracellular BRB maturation. We were not able to extend our studies beyond P14 in *Vglut1*^-/-^ mice, as the pups die around P15 ^71^. Notably, the vascular phenotypes, albeit opposite, were more severe in the *Vglut1*^-/-^ retina compared to the *Gnat1*^-/-^ retina. This is likely because glutamate release is abolished from both bipolar cells and photoreceptors (rods and cones) in the *Vglut1*^-/-^ retina ^12^. In contrast, only depolarized rod photoreceptors release excess glutamate in the *Gnat1*^-/-^ retina ^26, 27^. This may likely explain why the vascular phenotypes are less prominent in the P10 *Gnat1*^-/-^ retina, and possibly why the angiogenic phenotype (but not BRB phenotype) in the *Gnat1*^-/-^ retina normalizes to wild-type levels by P14.

How does extracellular glutamate trigger angiogenesis and BRB maturation in the developing retina? It is possible that the vascular phenotypes in *Vglut1*^-/-^ and *Gnat1*^-/-^ retinas could be due to alterations in either neuronal numbers which have been shown to affect retinal angiogenesis by titrating the level of available pro-angiogenic factor VEGF-A ^55^. However, we did not observe any significant differences in the numbers of neuronal subtypes between WT, *Vglut1*^-/-^ and *Gnat1*^-/-^ retinas. While there was a slight decrease in calbindin-labelled amacrine cells in the *Vglut1*^-/-^ retina, identified classes of amacrine cells, including cholinergic, GABAergic and dopaminergic neurons, were unaffected in these mutant retinas. These data suggest that angiogenesis and BRB phenotypes in *Vglut1*^-/-^ and *Gnat1*^-/-^ retinas are not caused by alterations in neuronal numbers. Metabolic demand of retinal neurons also modulates angiogenesis ^43, 67, 68^. However, the expression levels of HK2 and PKM2, two key glycolytic enzymes that are readouts of photoreceptor cell metabolism ^53^, were unaffected in both P10 *Vglut1*^-/-^ and *Gnat1*^-/-^ retinas compared to the WT, in spite of the prominent vascular phenotypes in these mutant retinas at this developmental stage. Similarly, retinal hypoxia as well as VEGF-A expression levels were also unaffected in both P10 *Vglut1*^-/-^ and *Gnat1*^-/-^ retinas compared to the WT. These data suggest that angiogenesis and BRB maturation abnormalities seen in these mutant retinas are not due to alterations in neuronal numbers and neuronal metabolic demand or VEGF-A signaling.

Our data demonstrate that *Norrin* mRNA expression and endothelial Norrin/β-catenin activity are downregulated in *Vglut1*^-/-^ retinas and upregulated in *Gnat1*^-/-^ retinas between P10-P14, and therefore are likely regulated by the extracellular glutamate concentration. Consistent with these findings our scRNA-seq analyses showed that expression levels of effectors of Norrin/β-catenin pathway [*Ctnnb1* and *Lef1*] are downregulated in *Vglut1*^-/-^ retinal ECs. However, *Norrin* mRNA is expressed in the retina before P10 ^20^, and perturbation of Norrin/β-catenin signaling delays superficial plexus angiogenesis as early as P5 ^20, 25^, well before the onset of glutamatergic neuronal activity. Therefore, early (P5-P8) *Norrin* mRNA expression in the retina, which is critical for superficial plexus angiogenesis, is likely regulated by a glutamate-independent mechanism ^48^. Our data in conjunction with a previous study ^48^ also suggest that *Norrin* mRNA is expressed by several cell types in the developing retina including Müller glia and INL neurons (horizontal cells and subsets of amacrine and bipolar cells). Using *ex vivo* organotypic cultures, we also show that glutamate can directly induce *Norrin* mRNA expression in the Müller glia and INL neurons, and this occurs via glutamate uptake by GLAST and GLT-1, two glutamate transporters expressed in Müller glia and INL neurons. We used a high concentration of TFB-TBOA (GLAST and GLT-1 inhibitor) in our study in order to ensure that glia- and neuron-mediated glutamate uptake is completely inhibited in our *ex vivo* culture system. Importantly, the presence of TFB-TBOA in the media in the absence of glutamate did not significantly alter *Norrin* mRNA expression compared to the retinas with neither glutamate nor TFB-TBOA in the media, confirming that TFB-TBOA does not affect *Norrin* mRNA expression in the retina. Future studies will elucidate the relative contribution of Müller glia- or INL neurons-derived Norrin, regulated by glutamate, for the deep plexus angiogenesis and paracellular BRB maturation. Importantly, pharmacological activation of endothelial Norrin/β-catenin signaling was sufficient to rescue both deep plexus angiogenesis and paracellular BRB integrity in the *Vglut1*^-/-^ retina, consistent with the idea that Norrin/β-catenin signaling functions downstream of extracellular glutamate to promote these processes. Our scRNA-seq data also show a slight downregulation of *Unc5b* in *Vglut1*^-/-^ retinal ECs. Unc5b has been shown to modulate Wnt/β-catenin signaling in CNS ECs ^72^, suggesting the possibility that the cross-talk between these two pathways may be affected in *Vglut1*^-/-^ retinal ECs. Interestingly, none of the downstream targets of VEGF-A signaling was affected in *Vglut1*^-/-^ retinal ECs at the transcriptome level. Thus, VEGF-A signaling pathway does not appear to be regulated by glutamate in the developing retina.

Taken together our data suggest a new mechanism by which glutamatergic neuronal activity regulate retinal angiogenesis and BRB maturation in the developing retina **(Figure 8L)**. We propose that extracellular glutamate, released either during spontaneous glutamatergic waves or light-dependent glutamatergic synaptic activity of photoreceptors from P8 onwards, signals to Müller glia and INL neurons. Glutamate signaling induces Norrin mRNA and protein expression and secretion by Müller glia and INL neurons. Norrin, in turn, activates endothelial Norrin/β-catenin signaling in ECs which promotes angiogenesis in the deep plexus and paracellular BRB maturation in both superficial and deep plexuses of the retina (**Figure 8L**). Our genetic data in *Vglut1*^-/-^ (no glutamate release into the synaptic cleft), *Gnat1*^-/-^ (excess glutamate release into the synaptic cleft) and *Chrnb*2^-/-^ (early onset of the glutamatergic waves), and the pharmacological rescue of *Vglut1*^-/-^ phenotypes by activation of endothelial Norrin/β-catenin signaling, are consistent with the proposed model that extracellular glutamate levels control deep plexus angiogenesis and barriergenesis. Although, the exact mechanisms by which glutamate mediates these effects are not fully clear, our single-cell RNA-seq data together with the published datasets ^48^ suggest that glutamate cannot act directly on ECs as they do not express either ionotropic or metabotropic glutamate receptor subunits or glutamate transporters. The published transcriptional data of the glutamate receptor and transporter expression ^48^, and our analysis of the response of the INL cells (neurons and Müller glia) to extracellular glutamate (i.e, changes in GS, GLAST, GLT-1 protein levels) in the *Vglut1*^-/-^ and *Gnat1*^-/-^ retinas, suggest that glutamate could directly act on Müller glia and INL neurons to induce *Norrin* mRNA expression. These cells express subsets of ionotropic or metabotropic glutamate receptors and glutamate transporters that make them exquisite sensors of extracellular glutamate levels. In addition, *Norrin* mRNA is predominantly expressed by the INL cells during vascular development and maturation. Future studies will address in more detail the molecular mechanisms by which glutamate affects Norrin expression in the INL cells and specific contributions of INL cell-derived Norrin to angiogenesis and barriergenesis.

Structural and functional abnormalities in neurons or glial cells precede vascular pathologies in multiple CNS neurological diseases with prominent vascular involvement, including those in the retina. For example, in diabetic retinopathy there is an early loss of neurovascular coupling, impaired response of photoreceptors to light, gradual neurodegeneration, gliosis and neuroinflammation before any observable vascular pathologies ^5–7^. Moreover, the early phases of either age-related macular degeneration, or retinitis pigmentosa, characterized by changes in synaptic activity and metabolic deficits in Müller glia ^73–75^, may also affect blood vessel function leading to neurodegeneration. Although vascular pathologies, including aberrant angiogenesis and BRB breakdown, are found in diabetic retinopathy and proliferative vitreoretinopathy that cause acquired adult blindness, as well as retinopathy of prematurity which leads to blindness in premature infants ^1, 76–78^, the molecular mechanisms that link neuronal or glial abnormalities with vascular deficits remain elusive. Our findings not only extend our understanding of the molecular mechanisms by which vascular development and homeostasis occurs in the CNS [reviewed in ^4^] including the role that neuronal activity plays in these processes both in the cortex ^79^ and retina ^14, 17, 18^, but also have important implications to identify putative interventions to treat CNS neurovascular pathologies, including those in the retina from diabetic retinopathy, retinitis pigmentosa to age-related macular degeneration.

## ACKNOWLEDGEMENTS

We thank Dr. Vilas Menon (Columbia University Irving Medical Center) for his consultation on computational analyses and Dr. Eduardo Solessio (Upstate Medical University, State University of New York) for his consultation about *Gnat1*^-/-^ retinal physiology. S.B., S.S., and D.A. are supported by grants from the National Eye Institute (R01EY033994 and K99EY033909), National Heart Lung and Blood Institute (R61HL159949, R33HL159949), National Institute of Aging (RF1AG078352), National Institute of Neurological Disorders and Stroke (R21 NS130265), the National Multiple Sclerosis Society (RG-1901-33218) and in part by a gift donation. W.J.B. and G.B. are supported by grants from the National Eye Institute (R01 EY033994 and R21 EY034696) and in part by funds from an unrestricted grant from Research to Prevent Blindness to the Department of Ophthalmology and Visual Sciences, Upstate Medical University.

## AUTHOR CONTRIBUTIONS

Conceptualization: S.B., W.J.B., D.A.; Methodology: S.B., D.A., G.B.; Software: S.S.; Validation: S.B., G.B; Formal analysis: S.B., S.S., G.B., P.A., D.J.; Investigation: S.B., S.S., G.B.; Resources: W.J.B., D.A.; Data curation: S.B., S.S., G.B.; Writing - original draft: S.B., D.A.; Writing - review & editing: S.B., S.S., G.B., W.J.B., D.A.; Visualization: S.B.; Supervision: W.J.B., D.A., Project administration: D.A., Funding acquisition: W.J.B., D.A.

## STAR ★ METHODS

### KEY RESOURCES TABLE

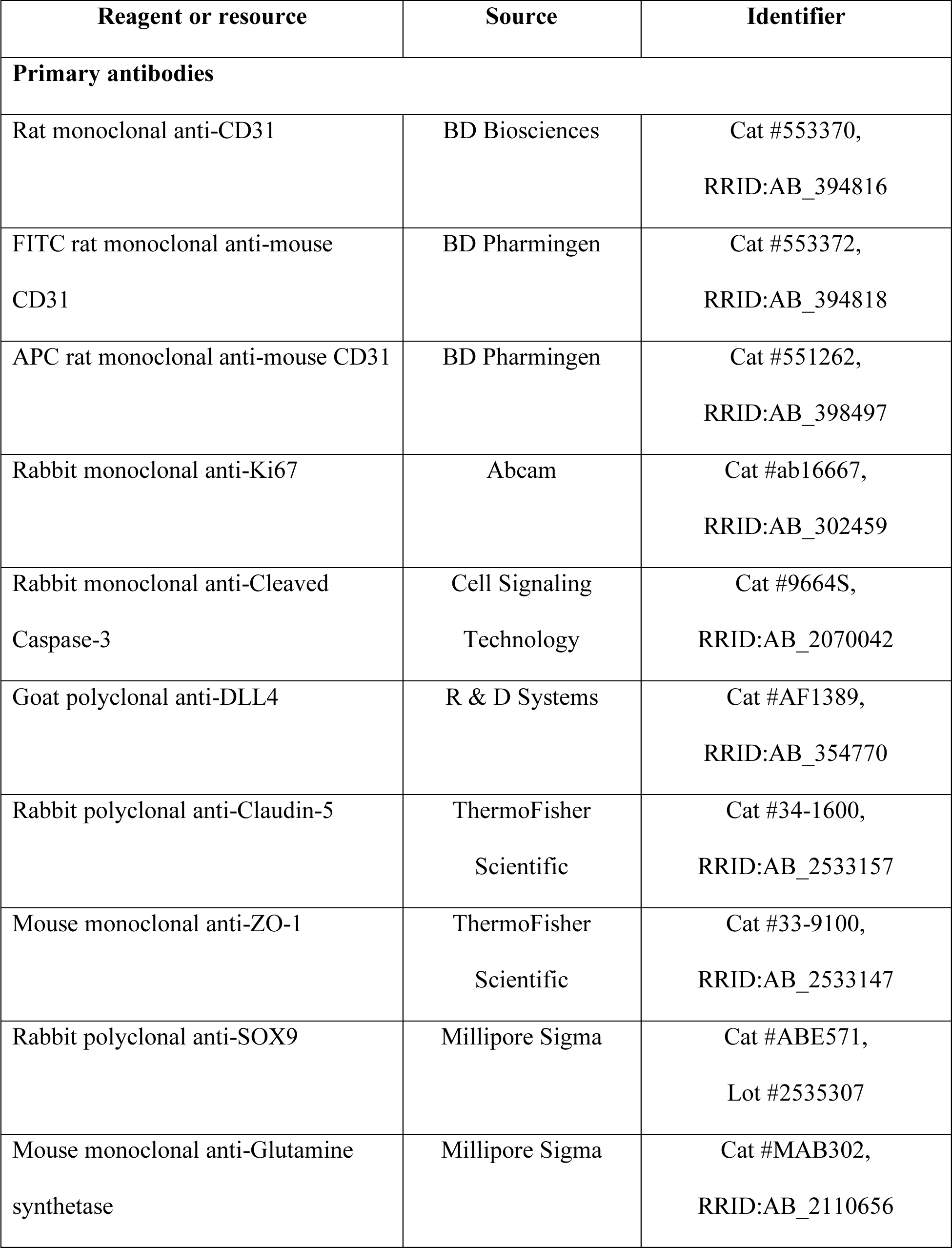
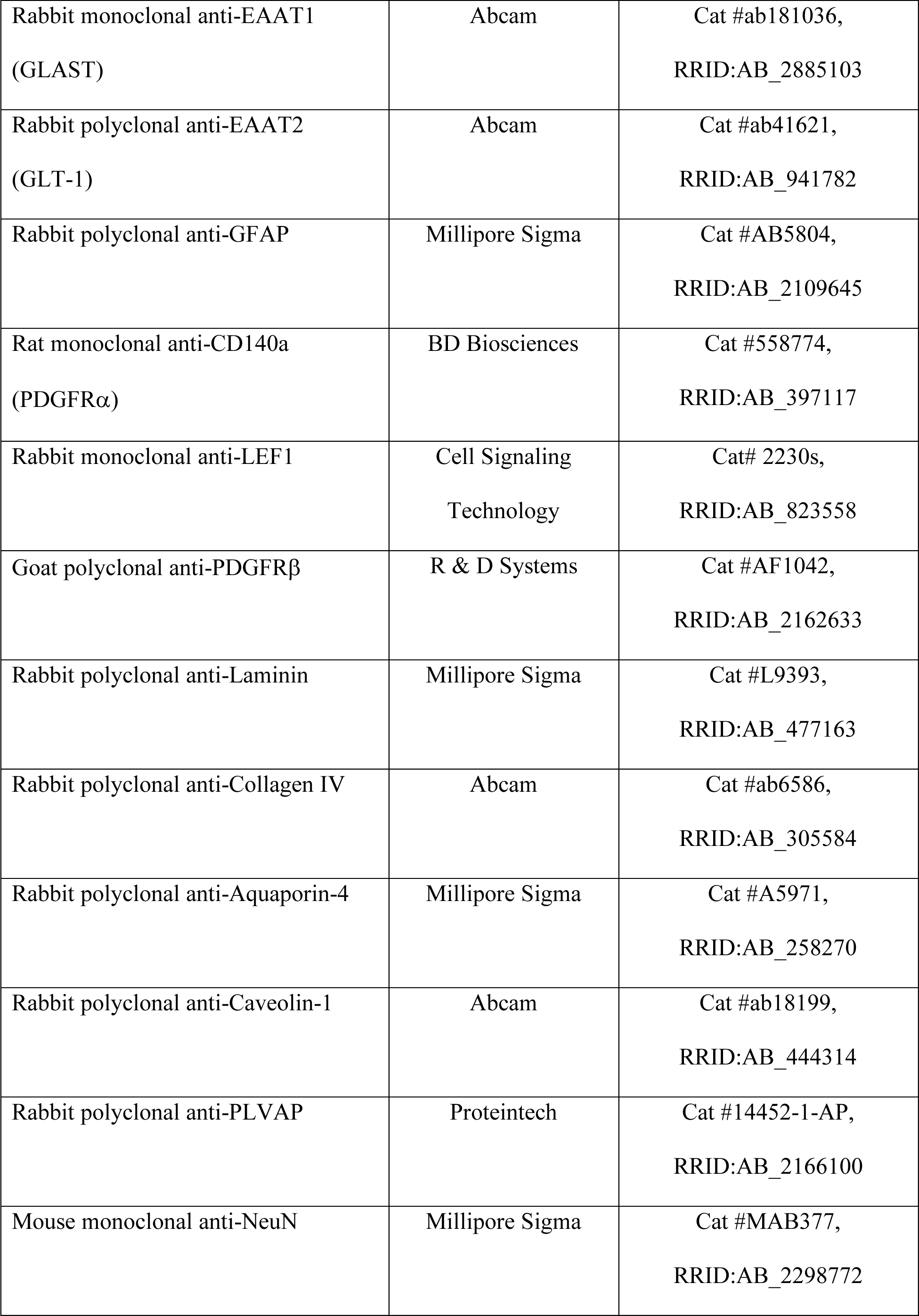
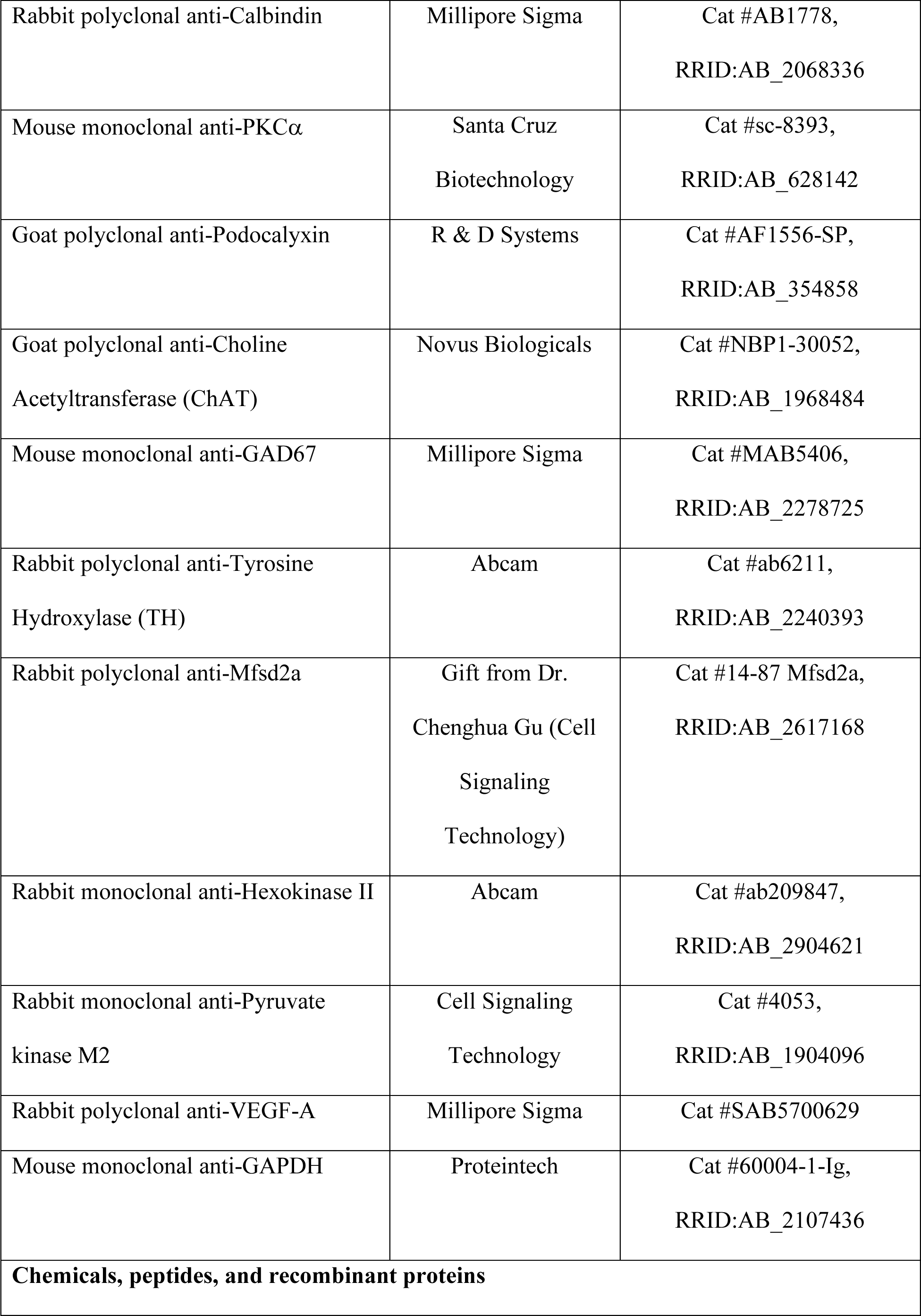
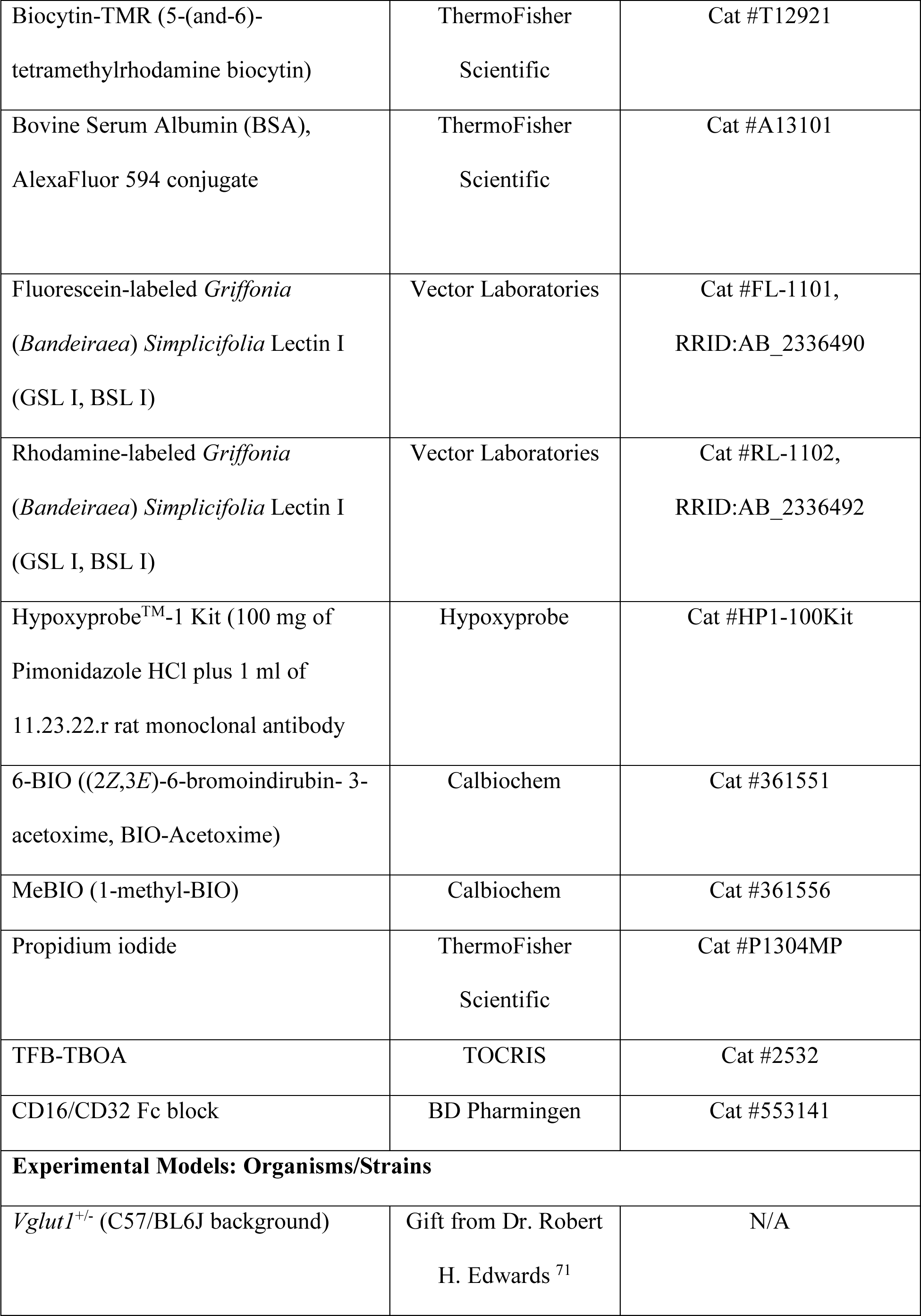
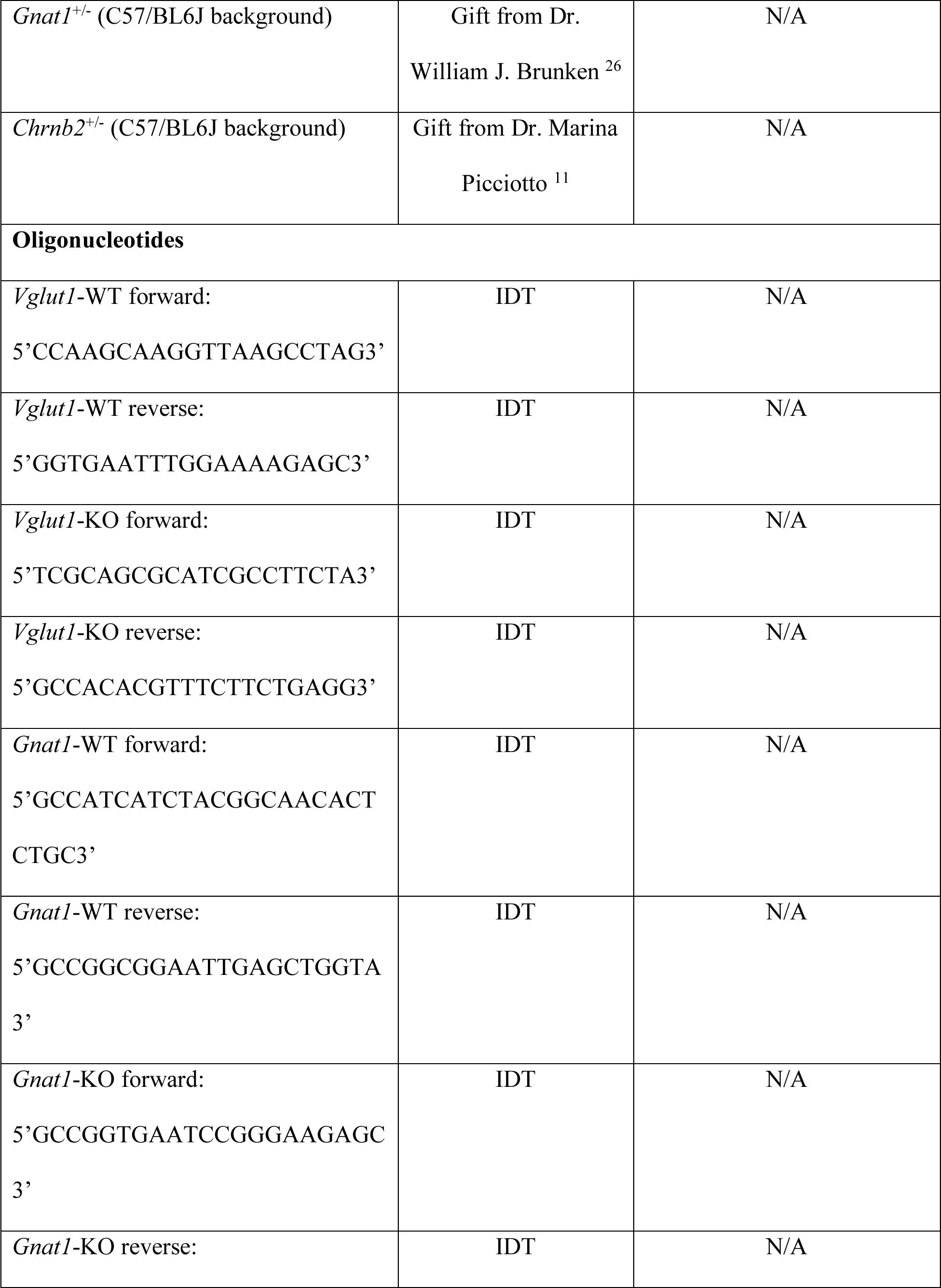
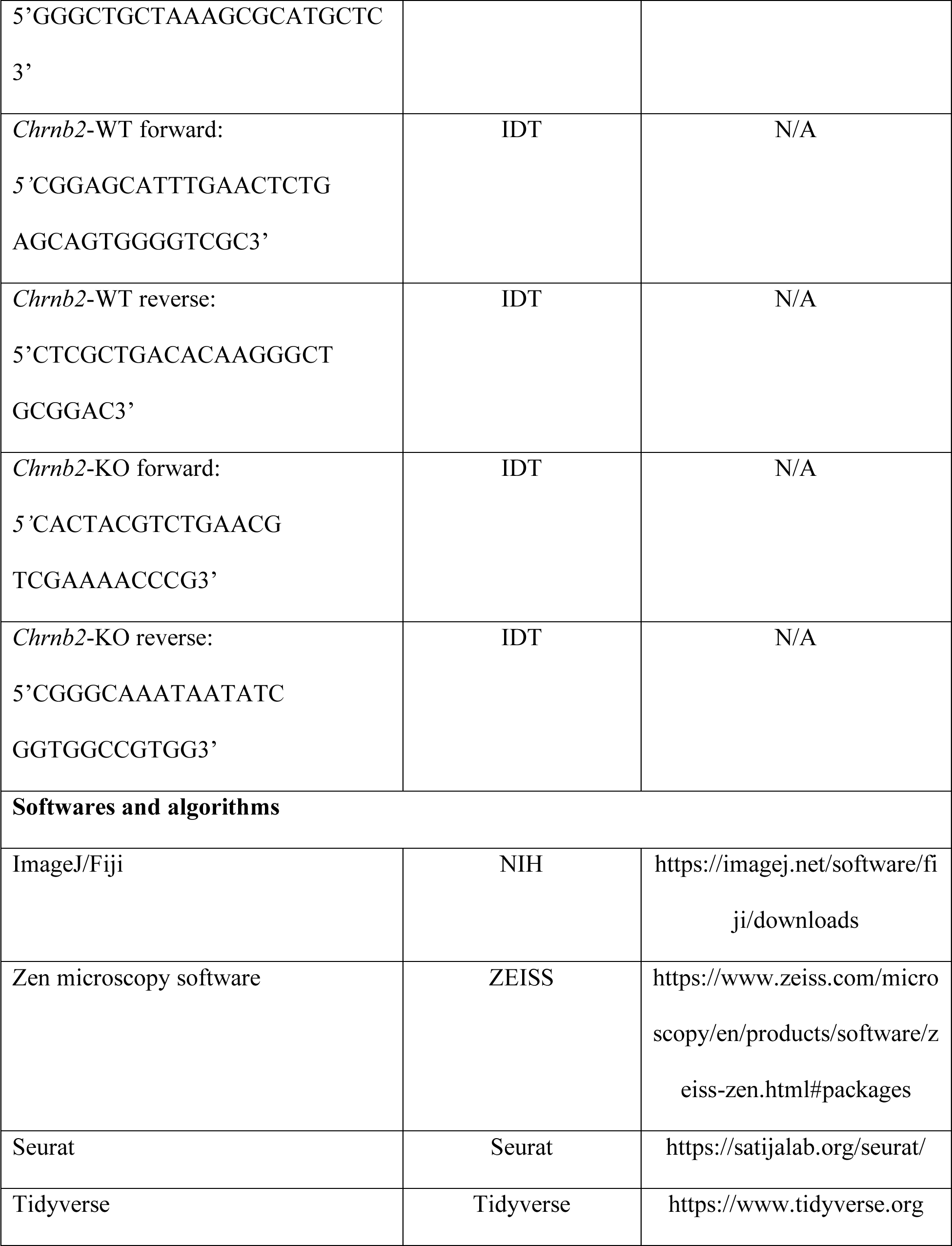
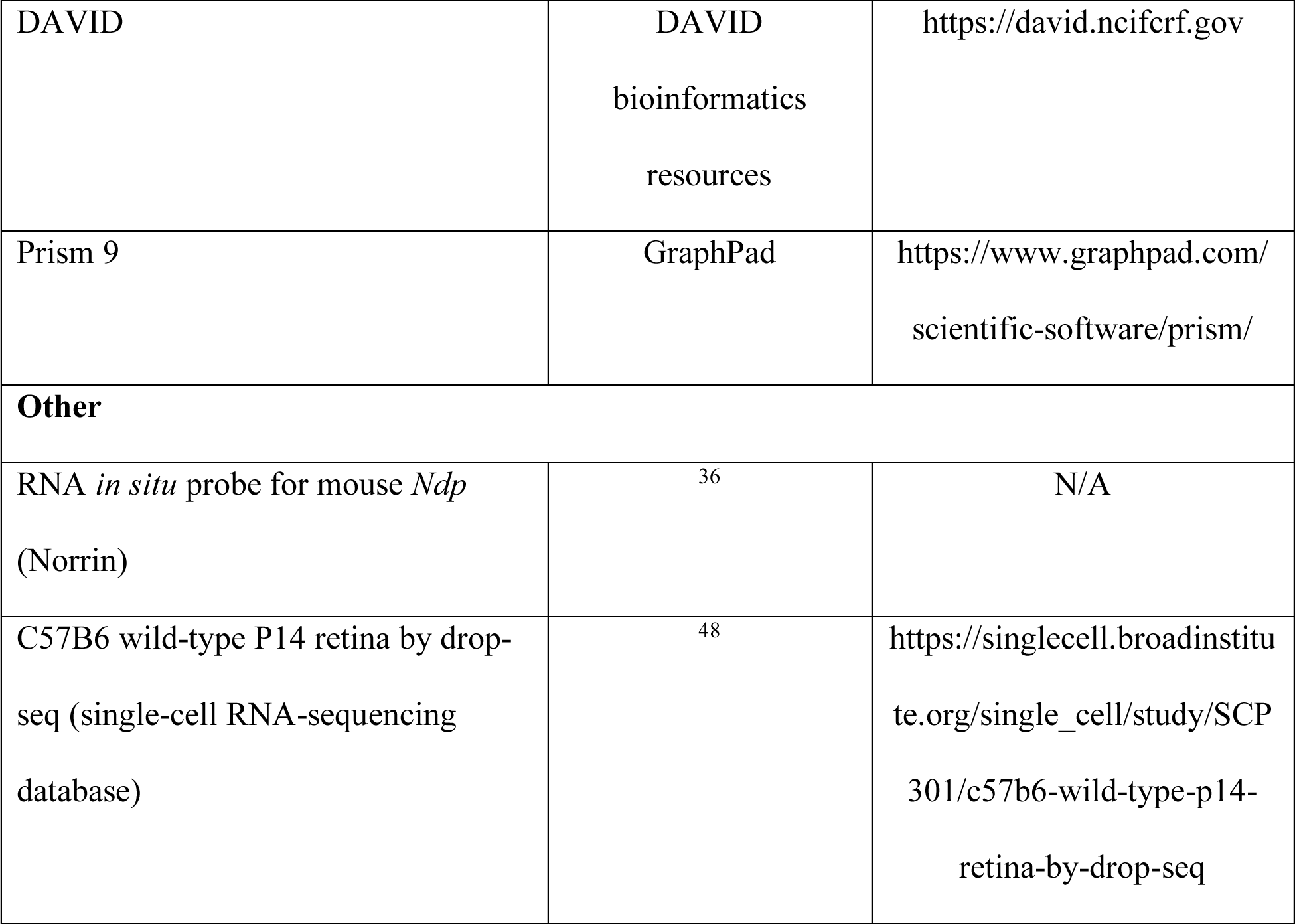

### RESOURCE AVAILABILITY

#### Lead contact

Further information and requests for resources and reagents should be directed to the lead contact, Dritan Agalliu.

#### Materials availability

This study did not generate new unique reagents.

#### Data and code availability

All data are available in the manuscript and the supplementary information. The codes used in this study have been previously published and are freely available (see Key Resources Table). Any additional information required to reanalyze the data reported in this study will be available upon request.

### EXPERIMENTAL MODEL AND SUBJECT DETAILS

#### Mice

All animal usage and protocols were approved by the Institutional Animal Care and Use Committee (IACUC) at Columbia University Irving Medical Center. Experimental procedures were optimized to minimize animal stress and the number of animals used for experiments. Mice were maintained on 12 light/12 dark cycle. Both sexes (males and females) were used for experiments and analyses. All mice were maintained in C57/BL6J background. The *in vivo* BRB permeability assays were performed either at P10 or P14. Generation of *Vglut1*^-/-^ mice was described before ^71^. These mice were bred as heterozygous and gave birth in Mendelian ratios. Age-matched littermate WT and *Vglut1*^-/-^ mice were used for experiments (*Vglut1*^-/-^ mice die around P15). Generation of *Gnat1*^-/-^ mice was described before ^26^. These mice were bred as heterozygous and gave birth in Mendelian ratios. Age-matched littermate WT and *Gnat1*^-/-^ mice were used for experiments. *Gnat1*^-/-^ mice are viable and fertile. Generation of *Chrnb2*^-/-^ mice was described before ^11^. These mice were bred as *Chrnb2*^+/-^ and gave birth in Mendelian ratios. Age-matched littermate WT and *Chrnb2*^-/-^ mice were used for experiments. *Chrnb2*^-/-^ mice are viable and fertile.

### METH1OD DETAILS

#### Immunofluorescence

Unless otherwise specified, isolated tissues (retina, brain and liver) were fixed in 4% PFA for 2 hours for immunostaining. The exceptions were retinal flat-mount staining for Laminin, in which cases retinas were fixed for 10 minutes in 2% PFA. For retinal flat-mounts, tissues were washed with 1X PBS following PFA fixation and blocked overnight in blocking buffer (5% goat or donkey serum, 0.3% Triton X-100 in 1× PBS) at 4°C. Samples were then incubated with primary antibodies in antibody-diluting solution (5% goat or donkey serum; 0.01% Triton X-100 in 1X PBS) for 48h at 4°C, washed, and incubated with secondary antibodies for 24 hours. For radial section staining, samples were washed in 1 × PBS, cryoprotected in 30% sucrose overnight, embedded in optimal cutting temperature compound (OCT; Tissue-Tek; Torrance, CA, USA), and sectioned at a thickness of 12 µm. Tissue sections were blocked for 2h at room temperature followed by primary antibody incubation overnight at 4°C. Following 1X PBS washes, sections were incubated with secondary antibodies for 4h at room temperature. Samples were imaged in room temperature using a Zeiss LSM700 confocal microscope [ZEISS 20X air (Objective Plan-Apochromat 20x/0.8 M27; numerical aperture: 0.8) or 40X water (Objective C-Apochromat 40x/1.2 W autocorr M27; numerical aperture: 1.2) objectives; ZEN image acquisition software] and analyzed using Fiji software.

#### Western blotting

Retinas were collected from mice after perfusion with PBS. Six retinas from three mice were combined for each sample. Collected tissue was homogenized in lysis buffer containing protease and phosphatase inhibitors using a Dounce homogenizer followed by sonication. Protein concentration was assayed by BCA (Bio-Rad) and resolved on SDS-PAGE. Protein levels in P10 retinas were measured by fluorescent western blot analysis [4-15% or 7.5% Mini-PROTEAN TGX gels (Bio-Rad)] and quantitation was performed using the LICOR system, as previously described. Fluorescent quantification of protein levels was performed using the Odyssey SA infrared imaging system. Values are displayed as protein levels corrected with GAPDH (loading control) and normalized to the WT values.

#### Antibodies

Primary antibodies used were rat anti-CD31 (BD Biosciences, Cat #553370, 1:250), rabbit anti-Ki67 (Abcam, Cat #ab16667, 1:500), rabbit anti-cleaved Caspase-3 (Cell Signaling Technology, Cat #9664S, 1:500), Goat anti-DLL4 (R & D Systems, Cat #AF1389, 1:200), rabbit anti-Claudin-5 (ThermoFisher Scientific, Cat #34-1600, 1:500), rabbit anti-SOX9 (Millipore Sigma, Cat #ABE571, 1:500), mouse anti-Glutamine Synthetase (Millipore Sigma, Cat #MAB302, 1:500), rabbit anti-EAAT1 (Abcam, Cat #ab181036, 1:250), rabbit anti-EAAT2 (Abcam, Cat #ab41621, 1:250), rabbit anti-GFAP (Millipore Sigma, Cat #AB5804, 1:500), rat anti-CD140a (BD Biosciences, Cat #558774, 1:100), rabbit anti-LEF1 (Cell Signaling Technology, Cat #2230s, 1:100), goat anti-PDGFRβ (R & D Systems, Cat #AF1042, 1:250), rabbit anti-Laminin (Millipore Sigma, Cat #L9393, 1:500), rabbit anti-Aquaporin-4 (Millipore Sigma, Cat #A5971, 1:250), rabbit anti-Caveolin-1 (Abcam, Cat #ab18199, 1:500), rabbit anti-PLVAP (Proteintech, Cat #14452-1-AP, 1:250), mouse anti-NeuN (Millipore Sigma, Cat #MAB377, 1:500), rabbit anti-Calbindin (Millipore Sigma, Cat #AB1778, 1:250), mouse anti-PKCα (Santa Cruz Biotechnology, Cat #sc-8393, 1:1000), rabbit anti-Mfsd2a (a gift from Dr. Chenghua Gu, 1:300), goat anti-Podocalyxin (R & D Systems, Cat #AF1556-SP, 1:500), goat anti-ChAT (Novus Biologicals, Cat #NBP1-30052, 1:100), mouse anti-GAD67 (Millipore Sigma, Cat #MAB5406, 1:2000), rabbit anti-Tyrosine Hydroxylase (Abcam, Cat #ab6211, 1:1000), rabbit anti-Hexokinase II (Abcam, Cat #ab209847, 1:1000), rabbit anti-Pyruvate Kinase M2 (Cell Signaling Technology, Cat #4053, 1:1000), rabbit anti-VEGF-A (Millipore Sigma, Cat #SAB5700629, 1:2000), mouse anti-GAPDH (Proteintech, Cat #60004-1-Ig, 1:50000), FITC rat anti-mouse CD31 (BD Pharmingen, Cat #553372, 1:200), and APC rat anti-mouse CD31 (BD Pharmingen, Cat #551262, 1:200). Secondary antibodies used were: goat anti-rat Alexa Fluor 594 (A11007), goat anti-rabbit Alexa Fluor 594 (A11012), goat anti-rat Alexa Fluor 488 (A11006), goat anti-rabbit Alexa Fluor 488 (A11034), donkey anti-rat Alexa Fluor 594 (A21209), donkey anti-rabbit Alexa Fluor 594 (A32754), donkey anti-rat Alexa Fluor 488 (A21208), donkey anti-rabbit Alexa Fluor 488 (A21206), donkey anti-rat Alexa Fluor 647 (A48272), donkey anti-rabbit Alexa Fluor 647 (A32795), donkey anti-goat Alexa Fluor 594 (A32758) and donkey anti-goat Alexa Fluor 488 (A11055) (all 1:250; ThermoFisher Scientific). IRDyes 680 and 800 (1:20000; LI-COR) were used as secondary antibodies for Western blot.

#### *In situ* hybridization

Both alkaline phosphatase and fluorescent *in situ* hybridization of fresh-frozen (in OCT) retinal sections using antisense mRNA probe for full-length mouse *Ndp* (Norrin) were performed as described before ^36^. The samples were imaged using either a ZEISS Axio Imager (alkaline phosphatase) or a ZEISS LSM700 confocal (fluorescent) microscope and analyzed using Fiji software.

#### Barrier permeability assays

Mice were anesthetized using isoflurane and injected with either 1% Biocytin-5(and-6-)-Tetramethylrhodamine (biocytin-TMR: ThermoFisher Scientific, Cat #T12921) or 1% Bovine Serum Albumin (BSA) - AlexaFluor 594 conjugate (albumin-Alexa594: ThermoFisher Scientific, Cat #A13101) into the tail veins. After 40 minutes in circulation, mice were re-anesthetized and perfused with 1X PBS via the cardiac route. Retinas and livers were isolated and fixed in 4% PFA for 2 hours prior to immunostaining with FITC-conjugated Lectin. Since blood vessels in the liver do not have a barrier, Biocytin-TMR or Albumin-594 fluorescence levels in the livers of corresponding animals were used as internal controls for measurement of biocytin-TMR and albumin-Alexa594 fluorescence levels in the retinas as described ^36, 37^.

#### Vessel perfusion assay

P14 pups were anesthetized using isoflurane and injected with 75 μl of Rhodamine-labeled Lectin (Vector Laboratories, Cat #RL-1102) into the left ventricles of the heart. Lectin was allowed to circulate for 5 minutes under anesthesia. After that, retinas were isolated and fixed in 4% PFA for 2 hours prior to immunostaining with vascular markers.

#### Hypoxia measurement

Hypoxyprobe staining was performed as per the vendor’s (Hypoxyprobe, Cat #HP1-100Kit) instructions. Briefly, the probe was dissolved in saline and administered into P10 pups by intraperitoneal injection at a concentration of 60 mg/kg body weight. After 90 minutes, the pups were anesthetized using isoflurane and perfused for 4 minutes with ice-cold sterile 1X PBS. After this, retinas were harvested, washed with 1X PBS following PFA fixation (4% PFA for 2 hours) and blocked overnight in blocking buffer at 4°C. Samples were then incubated with primary antibody (11.23.22.r) provided with the kit in antibody-diluting solution (1:50) for 48h at 4°C. Retinas were then washed with 1X PBS, and incubated with secondary donkey anti-rat Alexa594 antibody and Fluorescein-labeled Lectin for 24h.

#### Retinal organotypic culture

P10 WT pups were perfused with sterile 1X PBS under isoflurane-induced anesthesia, and eyeballs were enucleated. Eyeballs were dissected in sterile neurobasal media (ThermoFisher Scientific, Cat #21103049) supplemented with serum free B-27 (ThermoFisher Scientific, Cat #17504044). Corneas and lenses were removed from the eyeballs, and eyecups were placed in the upper chambers of sterile 12 mm Transwell with 0.4 μm Pore Polycarbonate Membrane Insert (Corning Life Sciences, Cat #3401). Eyecups were divided into four groups based on the media (neurobasal media + B-27) composition: 1) without glutamate, 2) with 0.1 mM glutamate, 3) without glutamate + 100 μm TFB-TBOA, 4) with 0.1 mM glutamate + 100 μm TFB-TBOA. The upper chambers of the transwell received 350 μl of the corresponding media, and the lower chambers of the transwell received 650 μl of the corresponding media. Under these conditions, the eyecups were placed in the sterile incubator for either 6 hours or 12 hours at 37° C and with 5% CO_2_. After 6 or 12 hours, the eyecups were removed from trans-well and fresh-frozen in OCT for future sectioning and fluorescent *in situ* hybridization.

#### Retinal endothelial cell isolation

P10 WT and *Vglut1*^-/-^ as well as P14 WT mice were anesthetized using isoflurane and perfused for 4 minutes with sterile 1X PBS. The eyes were enucleated and retinas were dissected in Earl’s balanced salt solution (EBSS). Retinas were dissociated in a single-cell suspension using a published protocol ^80^. Once the retinas were dissociated into single cells, it was resuspended in 200 ml of CD16/CD32 Fc block (BD Pharmingen, Cat #553141, 1:200 in FACS buffer) and incubated at room temperature for 15 minutes. The samples were washed with 2 ml of FACS buffer, with 100 ml kept aside for use as unstained and fluorescence minus one (FMO) controls. The cells were then resuspended in 200 ml of anti-mouse CD31 [either FITC rat anti-mouse CD31 (BD Pharmingen, Cat #553372, 1:200) or and APC rat anti-mouse CD31 (BD Pharmingen, Cat #551262), 1:200 in FACS buffer)] antibody to label ECs, covered in aluminum foil and incubated on ice for 60 mins. Following antibody labeling, the samples were washed twice with 2 ml of FACS buffer and resuspended in 400 ml of propidium iodide (Thermofisher, Cat. #P1304MP; 1/10,000 in FACS buffer). Gates for the surface stain were set using the unstained and FMO controls, and either CD31-APC^+^ or CD31-FITC^+^ ECs were sorted out using the BD FACSAria Flow Cytometer (Columbia Stem Cell Initiative Flow Cytometry Core) for single-cell RNA-sequencing.

#### Single-cell RNA-sequencing (scRNA-seq) analysis

Total of 1104 ECs for P10 WT retinas and 1046 ECs for P10 *Vglut1*^-/-^ retinas were analyzed for differential gene expression analyses (**Table S1**). The 10x Genomics Chromium platform ^81, 82^ was used to generate low-depth data (∼2000 genes/cell), and was processed using the Cell Ranger analysis pipeline to align reads and generate feature-barcode matrices. The Seurat R package ^83^ was implemented to read the output of the Cell Ranger pipeline and merge cells from all the samples (WT and mutant) into a single R object. The standard pre-processing workflow for scRNAseq data was carried out in Seurat, which consisted of the selection and filtration of cells based on quality control metrics (e.g., number of unique genes detected in each cell > 200, total number of molecules detected within a cell > 1,000 and < 50,000, percentage of reads within a cell that map to the mitochondrial genome < 20, etc.), data normalization and scaling, and the detection of highly variable features. Next, linear dimensionality reduction (PCA) was performed on the scaled data using the previously determined highly variable features, an alternative heuristic method (’Elbow plot’) was implemented to determine the ‘dimensionality’ of the dataset, and cells were clustered by applying the weighted shared nearest-neighbor graph-based clustering method using the first 20 PCs ^84^. Lastly, the 20 PCA dimensions were further reduced into two-dimensional space using the non-linear dimensional reduction technique, uniform manifold approximation and projection for dimension reduction (UMAP), to visualize and explore the scRNA-seq dataset and each of its unique EC clusters. All the ECs isolated from either genotype were plotted in the UMAP. Differential gene expression analyses were performed for all ECs from either genotype, and not on specific clusters. From the gene signatures that are uniquely present in mutant ECs compared to WT ECs, and validated by fluorescent in situ hybridization and/or immunofluorescence. Gene ontology (GO) enrichment analysis was performed using DAVID bioinformatics database to elucidate significantly up- and downregulated pathways in mutant ECs.

#### BIO and MeBIO treatment

The *in vivo* injections of 6-BIO ((2*Z*,3*E*)-6-bromoindirubin-3-acetoxime, BIO-Acetoxime; Calbiochem, Cat #361551) and control MeBIO (1-methyl-BIO; Calbiochem, Cat #361556) were performed as previously published ^64^. Pups were anesthetized with 1% isoflurane. Intraperitoneal injections of BIO and MeBIO were performed at a concentration of 5 mg/kg/day in sterile PBS from P6 to P8. Injected pups were re-anesthetized and perfused with 1X PBS to harvest tissues for analyses at P10.

### QUANTIFICATION AND STATISTICAL ANALYSIS

#### Image quantification

For retinal flat-mounts, each measurement was made in at least 3 quadrants per sample. The measurements were done separately in the central (near the optic nerve-head) and peripheral (near the edge) retina. For sections, at least three sections were imaged per sample. In each case, multiple representative images were acquired and averaged. To measure vascular area and branch point density, area positive for CD31 or Lectin was measured from each image in Fiji software (National Institutes of Health) using signal masking. Branch point numbers were normalized to unit area (1 mm^2^). To measure number of proliferative or apoptotic ECs, the number of Ki67^+^ or cleaved Caspase-3^+^ ECs, respectively, were normalized to unit CD31^+^ vascular area (1 mm^2^). The area of Biocytin-TMR leakage was done separately in the central and peripheral retina. The intensity of biocytin-TMR and albumin-Alexa594 outside of retinal vessels was measured as described ^36^. Biocytin-TMR and albumin-Alexa594 signal intensities in the retina and cerebella were normalized to those of livers. For vascular area positive for Claudin-5 and Caveolin-1, area positive for Lectin was selected from each image in Fiji software using signal masking, followed by measuring the percentage of Lectin^+^/Claudin-5^+^ or Lectin^+^/Caveolin-1^+^ in corresponding images. To measure the number of ganglion (NeuN^+^), amacrine, horizontal (Calbindin^+^) and bipolar cells (PKCα^+^), the number of cells were normalized to the length of either ganglion cell layer (GCL; for ganglion cells) or inner nuclear layer (INL; for amacrine, horizontal and bipolar cells). To measure the number of photoreceptors, the thickness of the outer nuclear layer (ONL) was quantified as a function of photoreceptor number. To measure the number of Müller glia, the number of Sox9^+^ cells were normalized to the length of INL. To measure the relative expression of GS and GLAST, the M.F.I were calculated for each field and data was presented as values normalized to the number of Sox9^+^ Müller glia. To measure *Ndp* expression, mean fluorescence intensity (M.F.I) of *Ndp* in the INL was calculated with histogram analysis in Fiji software and normalized to the WT values. For Lef1^+^ EC quantification, the number of Lef1^+^ ECs was normalized to unit CD31^+^ or Lectin^+^ vascular area (1 mm^2^). For western blots, values are displayed as target protein levels corrected with GAPDH (loading control) and normalized to the WT values using Odyssey SA infrared imaging system.

#### Statistical analysis

Retinas from 3-5 different animals (for *in vivo* experiments) were used for statistical analysis. For western blots, retinas from 3 animals/genotype were combined into one sample, and 4 samples/genotype was used for quantification. To test for statistical significance of any differences, unpaired Student’s *t* test or one-way ANOVA was performed using GraphPad Prism. A value of *P <* 0.05 was considered statistically significant. For scRNA-seq analysis, we considered transcripts with adjusted p value < 0.05, and >1.2-fold (log_2_fc > 0.25 and log_2_fc < −0.25) difference in expression as significant. For GO analysis, Gene sets with false discovery rate (FDR) < 0.01 were considered significant.

