## Supplementary Information for "Glutamatergic neuronal activity regulates angiogenesis and blood-retinal barrier maturation via Norrin/β-catenin signaling"

Glutamate, neuronal activity, angiogenesis, blood-retinal barrier, tight junctions, Norrin/β-catenin signaling

SUPPLEMENTARY INFORMATION

SUPPLEMENTARY FIGURES AND FIGURE LEGENDS

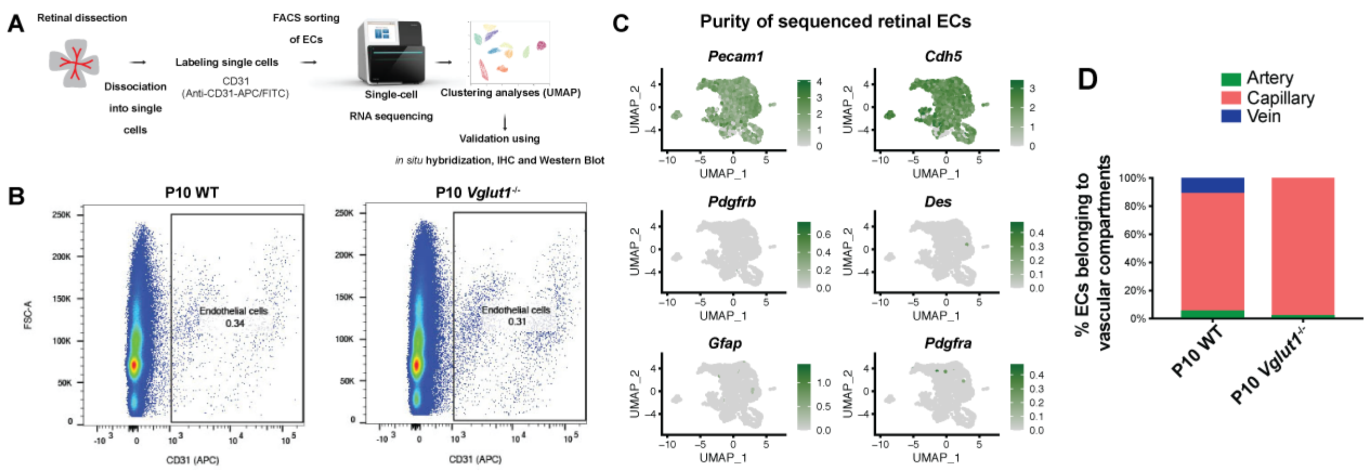


**Figure S1 (related to Figure 2). Isolation, single-cell RNA-sequencing, and validation of CD31$^{+}$ endothelial cells from WT and *Vglut1$^{-/-}$* retinas. A)** Schematic diagram of the experimental workflow for the isolation, transcriptomic profiling, and validation of endothelial cells (ECs) from P10 wild-type (WT) and *Vglut1$^{-/-}$* retinas. **B)** Fluorescence Cell Activating Sorting (FACS) plots show the gating strategy for sorting ECs from dissociated P10 WT and *Vglut1$^{-/-}$* retinas for single-cell RNA-sequencing (scRNA-seq) based on CD31 expression (x-axis), and the fraction of purified ECs from the total cell number in the retina for each genotype. **C)** Feature plots of *Pecam1* (CD31), *Cdh5* (EC markers), *Pdgfrb*, *Des* (pericyte markers), *Gfap* and *Pdgfra* (astrocyte markers) expression in the combined scRNA-seq data obtained from P10 WT and *Vglut1$^{-/-}$* retinal ECs and visualized with the Uniform Manifold Approximation and Projection (UMAP) plot. **D)** Stacked bar graphs showing the percentage of arterial (*Bmx$^{+}$*, green), venous (*Nr2f2$^{+}$*, blue) and capillary (*Mfsd2a$^{+}$*, red) ECs sequenced from each genotype. The majority of isolated retinal ECs from both genotypes were capillary ECs.

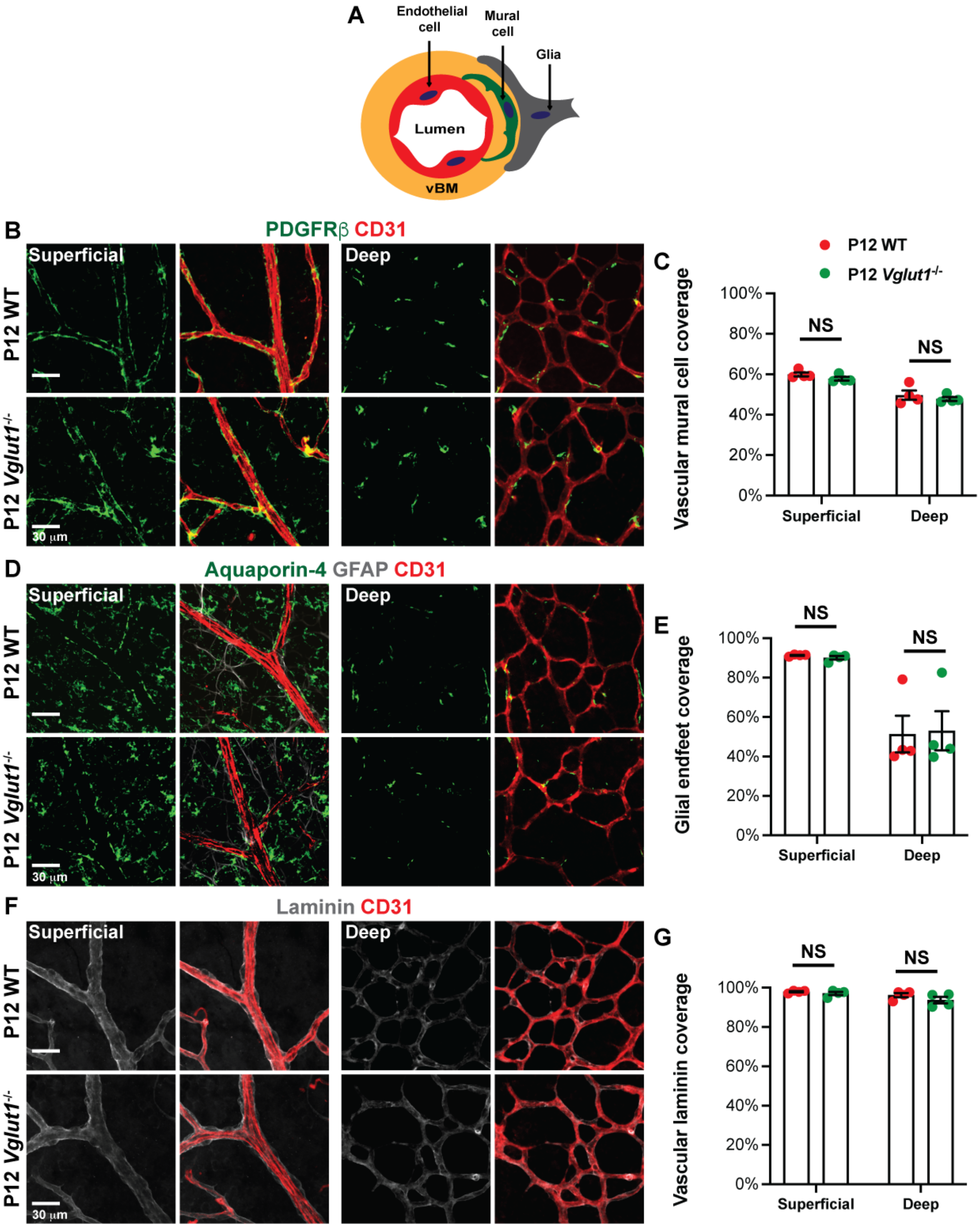


**Figure S2 (related to Figure 4). The organization of the neurovascular unit is intact in the *Vglut1*$^{-/-}$ retina. A)** Schematic diagram of the neurovascular unit (NVU) shows its cellular components and the vascular basement membrane (vBM). **B, D, F)** Representative images of superficial and deep plexuses in

P12 WT and *Vglut1*$^{-/-}$ retinal flat mounts labeled with PDGFRβ (**B,** green; mural cell marker), Aquaporin-4 (Aqp-4; **D,** green; glial endfeet marker) and Laminin (**F,** grey; vBM marker) along with CD31 (red; EC marker). **C, E, G)** Dotted bar graphs show the percentage of mural cell (**C**), glial endfeet (**E**) and vBM (**G**) coverage of retinal blood vessels. There is no difference in either mural cell, glial endfeet or vBM coverage of blood vessels in the superficial and deep plexuses between P12 WT and *Vglut1*$^{-/-}$ retinas (n = 4 mice / genotype). Scale bars = 62 μm. NS = not significant; Students t-test. Error bars: Mean ± S.E.M.

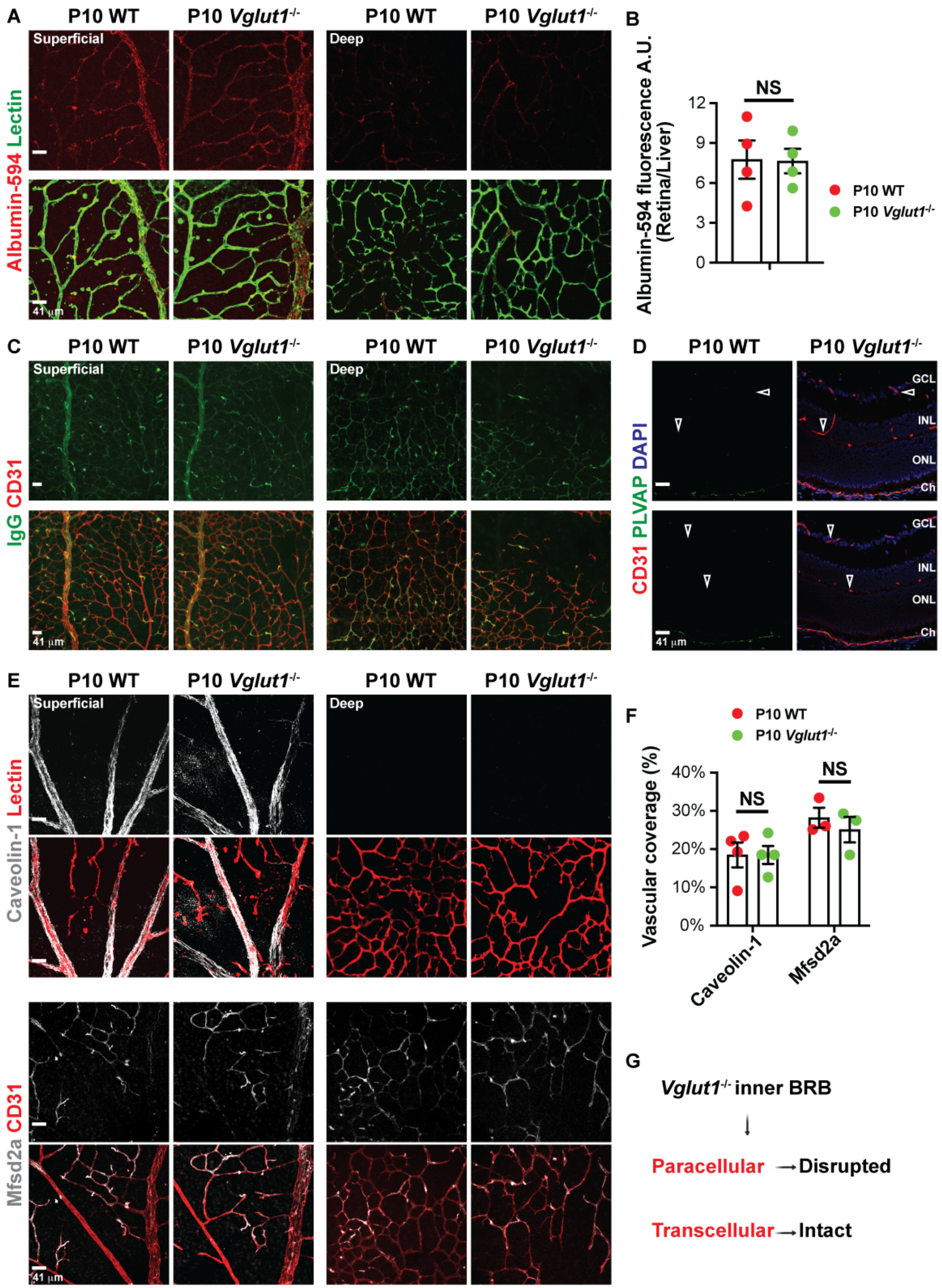


**Figure S3 (related to Figure 4). The maturation of the transcellular BRB occurs normally in the *Vglut1*$^{-/-}$ retina. A)** Representative images of the superficial and deep plexuses of P10 WT and *Vglut1*$^{-/-}$

retinal flat mounts labeled with Lectin (green) 40 minutes after intravenous injection of albumin-Alexa594 (red). Albumin-Alexa594 is confined almost entirely within blood vessels. **B)** Dotted bar graph of the fluorescence intensity of parenchymal albumin-Alexa594 in P10 WT and *Vglut1*$^{-/-}$ retinal flat mounts (n = 4 mice/genotype; Student's t-test). There is no difference between the two genotypes. **C)** Representative images of the superficial and deep plexuses of P10 WT and *Vglut1*$^{-/-}$ retinal flat mounts stained for CD31 (red) and endogenous serum IgG (green). Serum IgG is confined entirely within blood vessels. **D)** P10 WT and *Vglut1*$^{-/-}$ retinal sections were stained for PLVAP (green), CD31 (red) and DAPI (blue). PLVAP is present in the fenestrated choroidal blood vessels, but it is absent from retinal blood vessels (empty arrowheads) in both genotypes. **E)** Representative images of the superficial and deep plexuses of P10 WT and *Vglut1*$^{-/-}$ retinal flat mounts stained for Caveolin-1 (gray) or Mfsd2a (gray) proteins along with Lectin or CD31 (red). **F)** Dotted bar graph of Caveolin-1 and Mfsd2a coverage of blood vessels in P10 WT and *Vglut1*$^{-/-}$ retinas (n = 4 mice/genotype). There is no difference between the two genotypes. **G)** Summary of the paracellular and transcellular BRB phenotypes observed in the *Vglut1*$^{-/-}$ retina. GCL: ganglion cell layer, INL: inner nuclear layer, ONL: outer nuclear layer, Ch: choroid. Scale bars = 41 µm. NS = not significant; Students t-test. Error bars: Mean ± S.E.M.

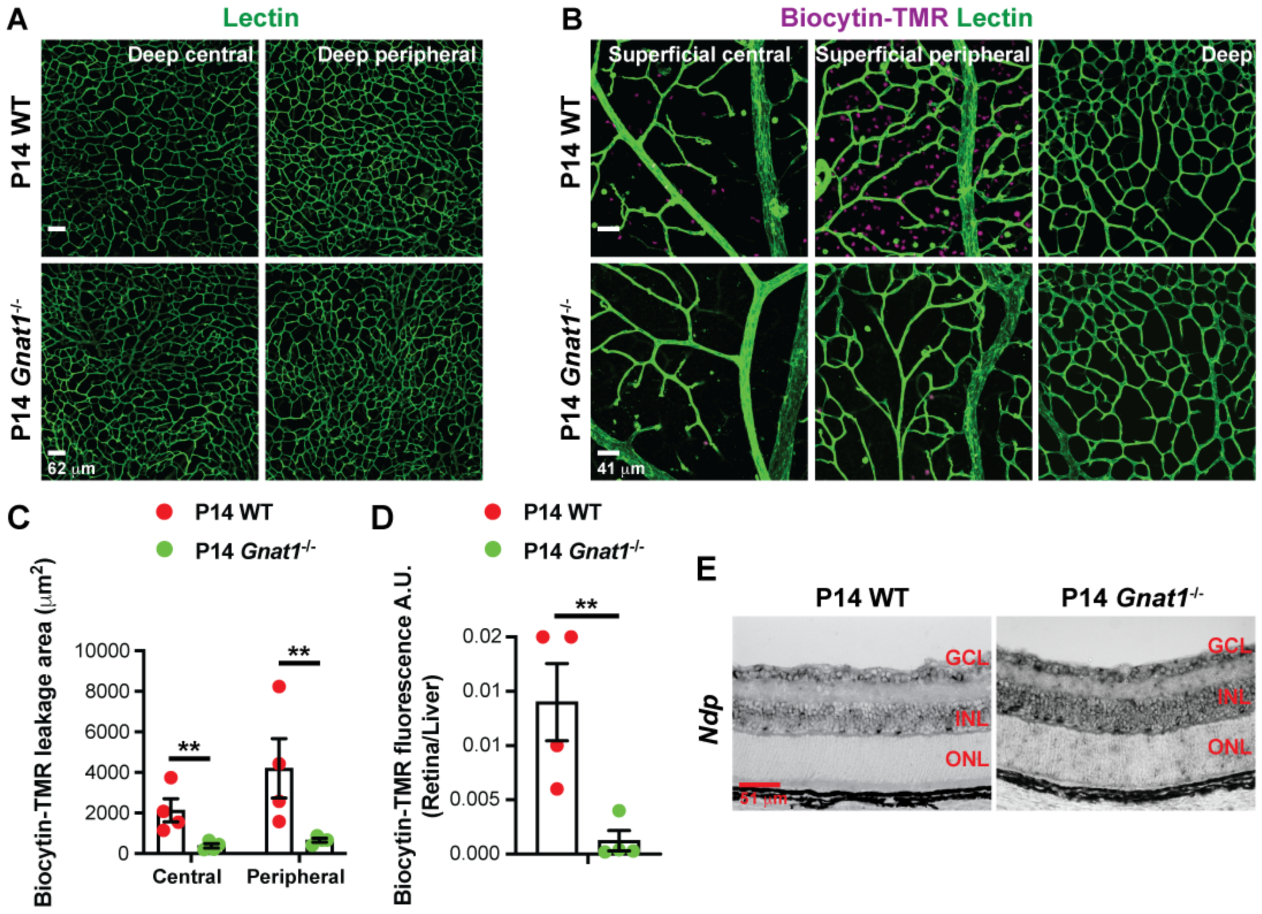


**Figure S4 (related to Figure 6). P14 *Gnat1*$^{-/-}$ retinas show precocious paracellular blood-retina barrier maturation and increased *Norrin* mRNA expression in the INL. A)** Representative images of the P14 WT and *Gnat1*$^{-/-}$ retinal deep plexuses stained for Lectin (green). **B)** Representative images of the superficial and deep plexuses from P14 WT and *Gnat1*$^{-/-}$ retinal flat mounts labeled with Lectin (green) 40 minutes after intravenous injection of biocytin-TMR (purple). **C, D)** Dotted bar graphs of the area (**C**) and fluorescence intensity (**D**) of parenchymal biocytin-TMR in P14 WT (red) and *Gnat1*$^{-/-}$ (green) retinal flat mounts (n = 4 mice / genotype). **E)** *In situ* hybridization for *Norrin* (*Ndp*) mRNA in P14 WT and *Gnat1*$^{-/-}$ retinas (n = 3 mice/genotype). *Norrin* mRNA signal intensity is increased in the INL layer of P14 *Gnat1*$^{-/-}$ compared to the WT retinas. GCL: ganglion cell layer, INL: inner nuclear layer, ONL: outer nuclear layer. Scale bars: **A** = 62 μm, **B** = 41 μm, E = 51 μm. Students t-test: ** p<0.02. Error bars: Mean ± S.E.M.

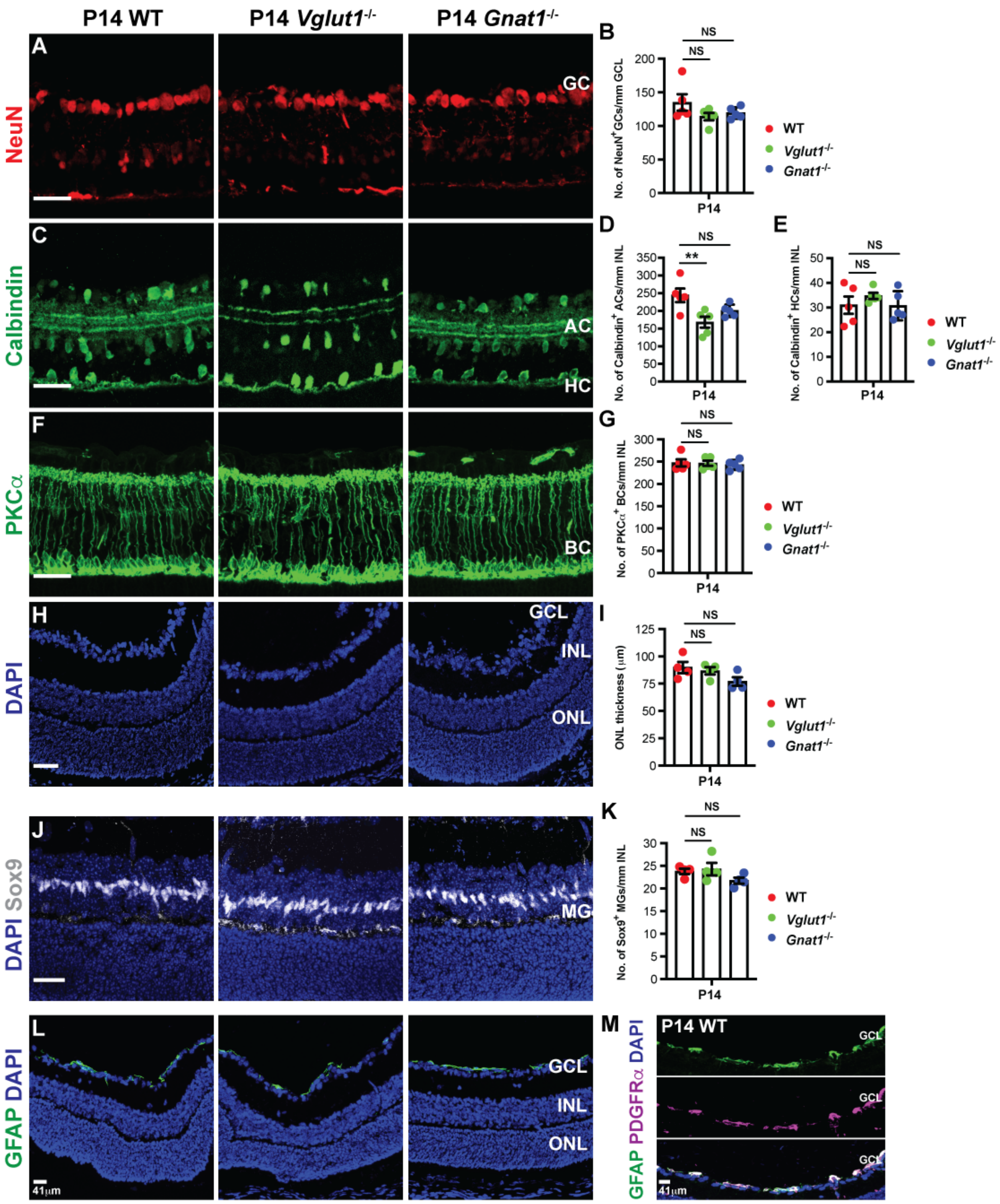


**Figure S5 (related to Figure 7). The number of neuronal subtypes and glial cells is largely unaffected in the *Vglut1*$^{-/-}$ and *Gnat1*$^{-/-}$ retinas. A, C, F, H)** P14 WT, *Vglut1*$^{-/-}$ and *Gnat1*$^{-/-}$ retinal sections were stained for either NeuN (**A**, red; ganglion cells), Calbindin (**C**, green; amacrine and horizontal cells), PKCα (**F**, green; rod bipolar cells), or DAPI (**H**). **B, D, E, G)** Dotted bar graph of the number of ganglion (**B**), amacrine (**D**), horizontal (**E**), and rod bipolar (**G**) cells per mm (section length) in P14 WT, *Vglut1*$^{-/-}$ and *Gnat1*$^{-/-}$ retinal sections (n = 5 mice / genotype). **I**) Dotted bar graph of the ONL thickness (readout of photoreceptor numbers) in P14 WT, *Vglut1*$^{-/-}$ and *Gnat1*$^{-/-}$ retinal sections (n = 4 mice /genotype). There is no difference among genotypes. **J, K)** P14 WT, *Vglut1*$^{-/-}$ and *Gnat1*$^{-/-}$ retinal sections were stained for Sox9 and DAPI (**J**), and the number of Sox9$^{+}$ Müller glia in the INL was quantified (**K**) for each genotype (n = 4 mice /genotype). There is no difference among groups. **L)** P14 WT, *Vglut1*$^{-/-}$ and *Gnat1*$^{-/-}$ retinal sections were stained for GFAP and DAPI. GFAP is only present in astrocytes in the GCL. **M)** P14 WT retinal sections were stained for GFAP and PDGFRα. GFAP is colocalized with PDGFRα in the GCL layer. GC: ganglion cell, AC: amacrine cell, HC: horizontal cell, BC: bipolar cell, MG: Müller glia, GCL: ganglion cell layer, INL: inner nuclear layer, ONL: outer nuclear layer. Scale bars = 41 μm. Statistical analyses were performed with one-way ANOVA with post-hoc Tukey; **p<0.02, NS = not significant. Error bars: Mean ± S.E.M.

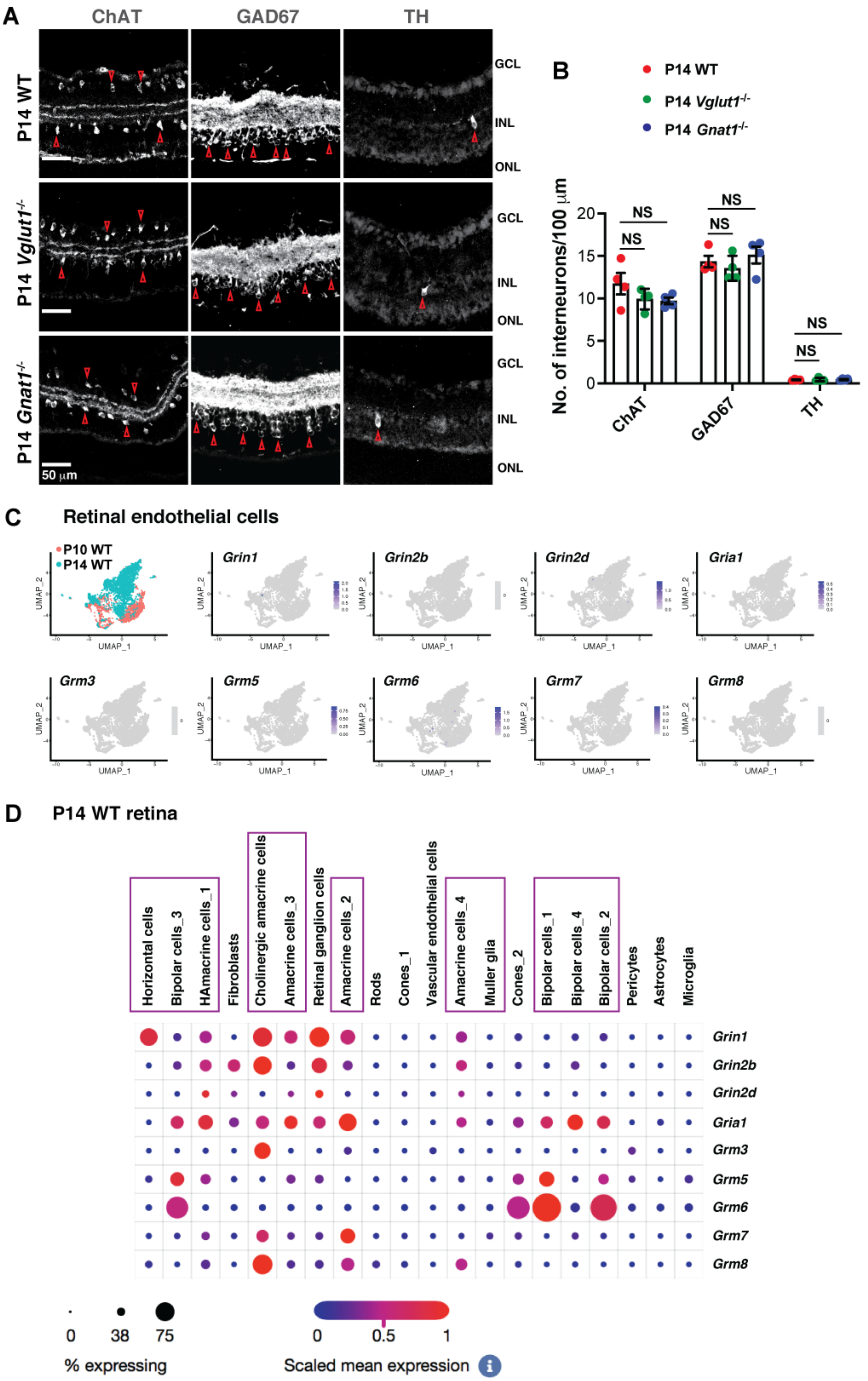
A
ChAT
GAD67
TH
P14 WT
P14 Vglut1-/-
P14 Gnat1-/-
GCL
INL
ONL
50 µm
B
P14 WT
P14 Vglut1-/-
P14 Gnat1-/-
No. of interneurons/100 µm
NS
ChAT
GAD67
TH
C
Retinal endothelial cells
P10 WT
P14 WT
Grin1
Grin2b
Grin2d
Gria1
Grm3
Grm5
Grm6
Grm7
Grm8
UMAP_1
UMAP_2
D
P14 WT retina
Horizontal cells
Bipolar cells_3
HAmacrine cells_1
Fibroblasts
Cholinergic amacrine cells
Amacrine cells_3
Retinal ganglion cells
Amacrine cells_2
Rods
Cones_1
Vascular endothelial cells
Amacrine cells_4
Muller glia
Cones_2
Bipolar cells_1
Bipolar cells_4
Bipolar cells_2
Pericytes
Astrocytes
Microglia
% expressing
Scaled mean expression

**Figure S6 (related to Figure 7). Glutamate receptors are not expressed in ECs during retinal vascular development. A)** P14 WT, *Vglut1*$^{-/-}$ and *Gnat1*$^{-/-}$ retinal sections were stained for ChAT (cholinergic neurons), GAD67 (GABAergic neurons) and Tyrosine Hydroxylase (TH; dopaminergic neurons). Empty red arrowheads point to representative neurons for each marker. GCL: ganglion cell layer, INL: inner nuclear layer, ONL: outer nuclear layer. **B)** Dotted bar graph of the number of ChAT$^{+}$, GAD67$^{+}$ and TH$^{+}$ neurons per 100 μm (section length) in P14 WT, *Vglut1*$^{-/-}$ and *Gnat1*$^{-/-}$ retinal sections (n = 4 mice / genotype). Scale bars = 50 μm. Statistical analyses were performed with one-way ANOVA; NS = not significant. Error bars: Mean ± S.E.M. **C)** UMAP projection of P10 and P14 WT retinal ECs that were FACS-sorted from the retina after enzymatic dissociation and processed for single-cell RNA-sequencing (scRNA-seq). Feature plots for ionotropic (*Grin1*, *Grin2b*, *Grin2d*, *Gria1*) and metabotropic (*Grm3*, *Grm5*, *Grm6*, *Grm7*, *Grm8*) glutamate receptor subunits mRNA expression superimposed in the UMAP projection. Metabotropic and ionotropic subunit receptor mRNAs are not expressed in either P10 or P14 retinal ECs. **D)** Dot plot of the expression profiles of the same glutamate receptors in the P14 WT retina from a published and available scRNA-seq database (please see text for more details). The dot size indicates the percent of the population expressing each marker; the color scale indicates the average level of gene expression. Metabotropic and ionotropic subunit receptor mRNAs are expressed at high levels in the inner nuclear layer (INL) neurons and Müller glia (marked with purple boxes).

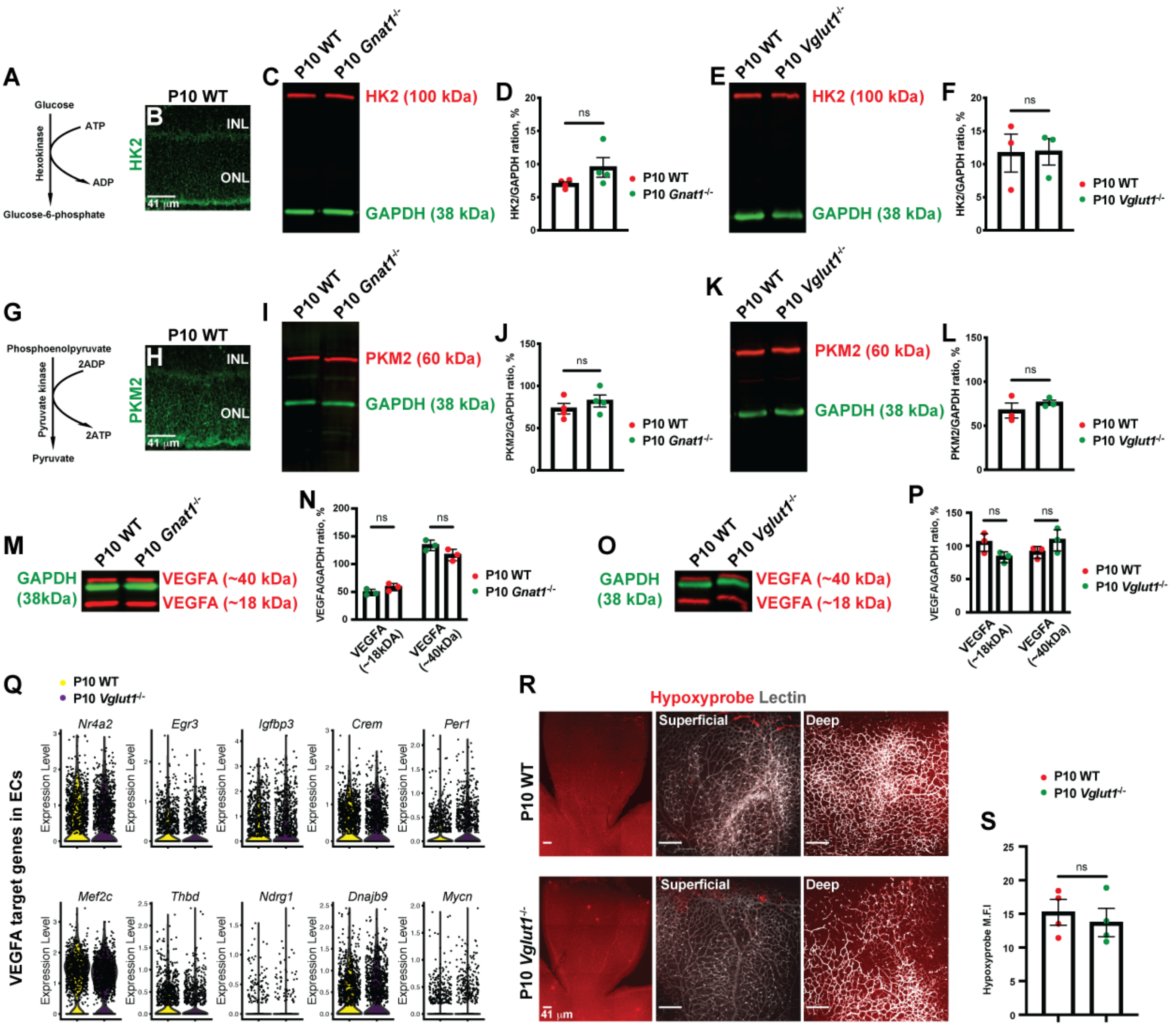


**Figure S7 (related to Figure 7). There are no differences in the metabolic demand and VEGF-A signaling among WT, *Vglut1*$^{-/-}$ and *Gnat1*$^{-/-}$ retinas. A)** Schematic diagram of the first step of glycolysis mediated by the Hexokinase (HK). **B)** P10 WT retinal section stained for HK2 (green) shows expression restricted primarily to the photoreceptors. **C, D)** Western blot and quantification for HK2 (red; 100 kDa) and GAPDH (green; 38kDa; loading control) proteins from P10 WT and *Gnat1*$^{-/-}$ retinal lysates (n = 4 mice/genotype). There is no difference between the genotypes. **E, F)** Western blot and quantification for HK2 (red; 100 kDa) and GAPDH (green; 38kDa; loading control) proteins from P10 WT and *Vglut1*$^{-/-}$ retinal lysates (n = 3 mice/genotype). There is no difference between the genotypes. **G)** Schematic diagram of the last step of glycolysis mediated by Pyruvate Kinase (PK). **H)** P10 WT retinal section stained for PKM2 (green) shows expression mainly in the photoreceptors. **I, J)** Western blot and quantification for

PKM2 (red; 62 kDa) and GAPDH (green; 38kDa; loading control) proteins from P10 WT and *Gnat1*$^{-/-}$ retinal lysates (n = 4 mice/genotype). There is no difference between genotypes. **K, L)** Western blot and quantification for PKM2 (red; 100 kDa) and GAPDH (green; 38kDa; loading control) proteins from P10 WT and *Vglut1*$^{-/-}$ retinal lysates (n = 3-4 replicates/genotype). There is no difference between genotypes. **M, N)** Western blot and quantification for VEGF-A (red; 18 kDa and 40 kDa) and GAPDH (green; 38kDa; loading control) proteins from P10 WT and *Gnat1*$^{-/-}$ retinal lysates (n = 3 mice/genotype). There is no difference between the two genotypes. **O, P)** Western blot and quantification for VEGF-A (red; 18 kDa and 40 kDa) and GAPDH (green; 38kDa; loading control) proteins from P10 WT and *Vglut1*$^{-/-}$ retinal lysates (n = 3 mice/genotype). There is no difference between the two genotypes. **Q)** Violin plots of transcript levels for VEGF-A target genes in P10 WT and *Vglut1*$^{-/-}$ retinal ECs from scRNA-seq. There is no difference between the two genotypes. **R)** Hypoxyprobe (Pimonidazole; red) and Lectin (gray) labeling of P10 WT and *Vglut1*$^{-/-}$ retinal flat mounts. **S**) Quantification of hypoxyprobe labeling intensity in P10 WT and *Vglut1*$^{-/-}$ retinas (n = 4 mice / genotype). Scale bars = 41 μm. Statistical tests were performed with the Students t-test: NS = not significant. Error bars: Mean ± S.E.M.

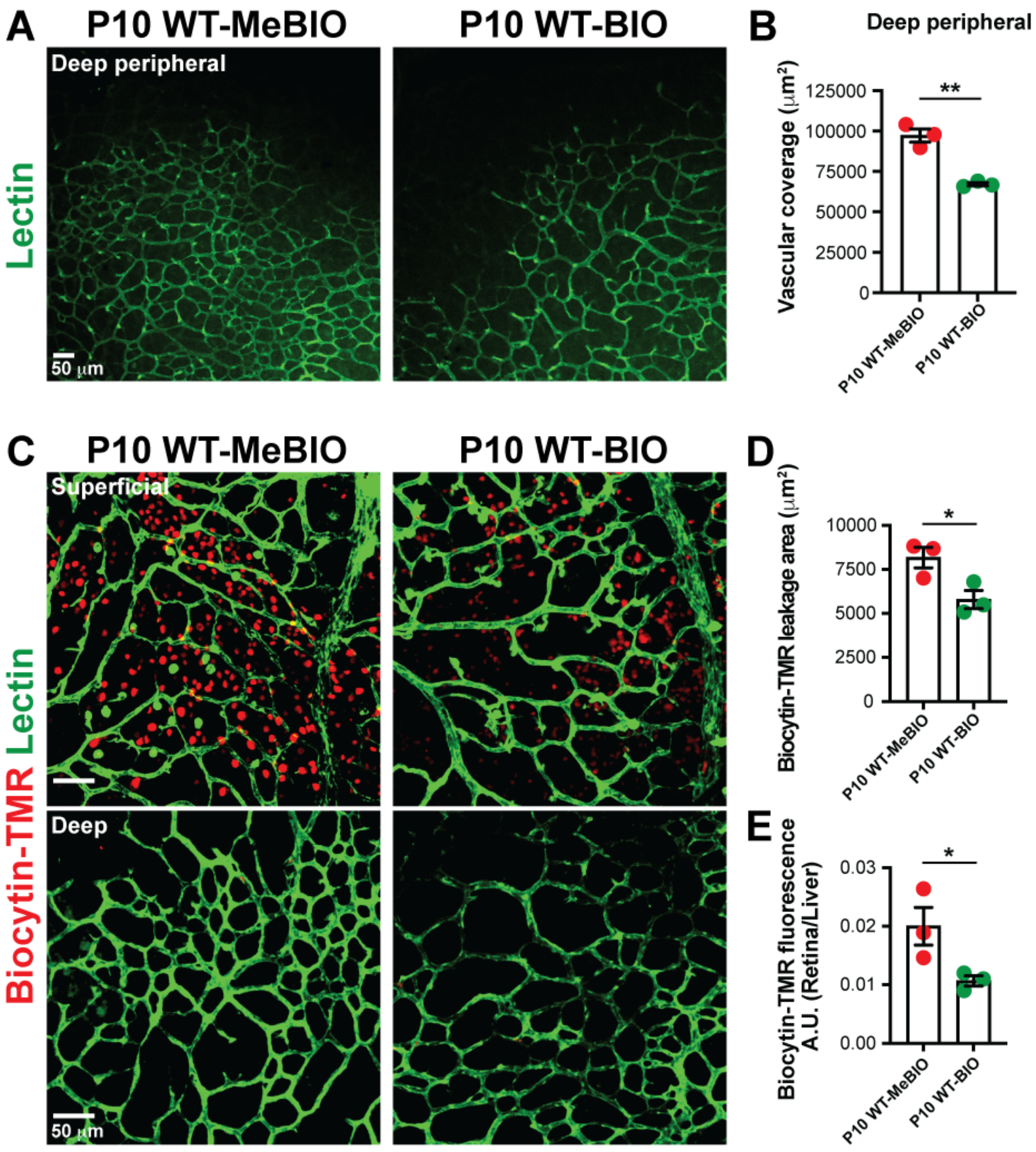


**Figure S8 (related to Figure 8). BIO treatment precociously matures paracellular BRB, but decreases deep plexus vascular coverage in wild-type retinas. A, B)** MeBIO- or BIO-treated P10 WT retinal flat-mounts were stained (**A**) with Lectin (green), and the vascular coverage of the peripheral deep plexus for each treatment was quantified (**B**) for each treatment (n = 3 mice / group). **C)** Representative images of flat-mount MeBIO- or BIO-treated P10 WT retinas 40 minutes after intravenous injection of biocytin-TMR (red) and stained with Lectin (green). **D, E)** Dotted bar graphs show quantification of biocytin-TMR leakage area (**D**) and fluorescence intensity (**E**) (n = 3 mice / group). Students t-test: * $p<0.05$, ** $p<0.02$. Error bars: Mean ± S.E.M. Scale bars = 50 μm.

### SUPPLEMENTARY DATASHEET LEGEND

**Tables S1-S7:** Batch structure **(Table S1)** of the number of ECs analyzed for the single-cell RNA-sequencing of the P10 *Vglut1*$^{-/-}$ retina compared to the P10 WT retina. List of all significantly differentially expressed EC genes **(Table S2)** between the P10 *Vglut1*$^{-/-}$ and WT retinal ECs. Gene ontology (GO) analyses of significantly downregulated **(Table S3)** and upregulated **(Table S4)** EC genes in the P10 *Vglut1*$^{-/-}$ retina compared to the P10 WT retina. Significantly downregulated and upregulated genes related to angiogenesis **(Table S5)**, EC tip cell **(Table S6)** and blood-retinal barrier (BRB) **(Table S7)** in the P10 *Vglut1*$^{-/-}$ retina compared to the P10 WT retina.
